# Metabolic adaption to extracellular pyruvate triggers biofilm formation in *Clostridioides difficile*

**DOI:** 10.1101/2021.01.23.427917

**Authors:** Yannick D.N. Tremblay, Benjamin A.R. Durand, Audrey Hamiot, Isabelle Martin-Verstraete, Marine Oberkampf, Marc Monot, Bruno Dupuy

## Abstract

*Clostridioides difficile* infections are associated with gut microbiome dysbiosis and are the leading cause of hospital acquired diarrhoea. The infectious process is strongly influenced by the microbiota and successful infection relies on the absence of specific microbiota-produced metabolites. Deoxycholic acid (DOC) and short chain fatty acids are microbiota-produced metabolites that limit the growth of *C. difficile* and protect the host against this infection. In a previous study, we showed that DOC causes *C. difficile* to form strongly adherent biofilms after 48 h. Here, our objectives were to identify and characterize key molecules and events required for biofilm formation in the presence of DOC. We applied time-course transcriptomics and genetics to identify sigma factors, metabolic processes and type IV pili that drive biofilm formation. These analyses revealed that extracellular pyruvate induces biofilm formation in the presence of DOC. In the absence of DOC, pyruvate supplementation was sufficient to induce biofilm formation in a process that was dependent on pyruvate uptake by the membrane protein CstA. In the context of the human gut, microbiota-generated pyruvate is a metabolite that limits pathogen colonization. Taken together our results suggest that pyruvate-induced biofilm formation might act as a key process driving *C. difficile* persistence in the gut.

## Introduction

*Clostridioides difficile*, formerly known as *Clostridium difficile*, causes infections associated with gut microbiome dysbiosis and is the leading cause of nosocomial diarrhea and colitis following antibiotic therapy (Crobach et al., 2018). While infections are typically associated with dysbiosis, recent epidemiological studies indicate that 5-15% of the population asymptomatically carry *C. difficile* despite having a healthy microbiota (Crobach et al., 2018). There is also increasing evidence that *C. difficile* causes community acquired infections and is a zoonotic pathogen. Pets and farm animals asymptomatically carry this pathogen and, as a result it is detected in retail meat. Based on these findings, *C. difficile* can now be viewed as a quintessential “One Health” pathogen (Lim et al. 2020).

The *C. difficile* infectious cycle depends on the ability of this anaerobic Gram-positive rod to sporulate. Once ingested from the surrounding environment or food, spores germinate in the ileum and then vegetative cells colonize the caecum (Crobach et al., 2018; Lim et al. 2020). Successful colonization relies on the disruption of the host’s microbiota (Abbas and Zackular, 2020; Ghimire et al. 2020; Girinathan et al., 2020; Pereira et al., 2020). The microbiota protects against *C. difficile* infection by producing metabolites including deoxycholic acid (DOC) and short chain fatty acids (SCFA) (Buffie et al., 2015; Studer et al. 2016; McDonald et al., 2018; Sobach et al. 2018; Seekatz et al., 2018). During dysbiosis specific members of the microbiota are missing, which results in altered SFCA and DOC production, thus allowing *C. difficile* to grow unimpeded (Abbas and Zackular, 2020). Following successful treatment of *C. difficile* infections (*via* antibiotic therapy or fecal transplant), the production of protective metabolites by the microbiota is slowly restored. Yet, relapses occur in more than 30% of patients following their first *C. difficile* infection and this rate increases to 50% after an initial relapse (Crobach et al., 2018). The exact cause of relapse has not been fully elucidated and represents a major challenge for managing *C. difficile* infections. In 40% of relapse cases, patients are infected with the same strain that caused the initial infection, suggesting *C. difficile* persists in the gastrointestinal tract (Crobach et al., 2018).

Persistence was initially associated with sporulation during antibiotic treatment, followed by germination after treatment. Evidence for this hypothesis was based on the inability of a non-sporulating *spo0A*-inactivated strain to persist and cause relapse in a murine infection model (Deakin et al. 2012). Inactivation of *spo0A* has pleiotropic effects influencing flagellar motility, metabolism, and biofilm-formation (Pettit et al. 2014; Dawson et al. 2012). This suggests that factors other than sporulation may also contribute to *C. difficile* persistence. We hypothesize that *C. difficile* forms multi-species biofilms in the gut which drives persistence in the presence of a normal microbiota (Donelli, et al, 2012; Crowther et al. 2014; Semenyuk et al, 2015). In support of this hypothesis, recent data from our laboratory showed that co-culture of *C. difficile* with *Clostridium scindens*, a bacterium that converts primary bile salts to secondary bile salts, promotes dual-species biofilm formation in the presence of cholate (Dubois et al. 2019).

Typically, biofilms are defined as community of bacterial cells encased in self-produced polymeric matrix (Hall-Stoodley et al., 2004). This matrix is usually composed of any combination of proteins, extracellular DNA, or exopolysaccharides and matrix composition will vary from species to species. Biofilms provide a microenvironment that decrease the susceptibility of bacteria towards different environmental stressors, including antimicrobial compounds, to promote bacterial persistence. In *C. difficile*, biofilm formation is mediated by several surface structures (e.g. pili and the S-layer), environmental triggers (e.g. DOC and sub-MIC of antibiotics), quorum sensing (e.g. *luxS*), in addition to other determinants (e.g. c-di-GMP) (Ðapa et al., 2013; Soutourina et al., 2013; Boudry et al., 2014; Pantaléon et al., 2015; Walter et al., 2015; Maldarelli et al., 2016; Vuotto et al. 2016).

The aim of this study is to identify key factors that contribute to biofilm formation by *C. difficile* in response to DOC. We used time-course transcriptomics and genetic techniques to identify sigma factors, metabolic processes and adhesins that drive biofilm formation. We then demonstrate that extracellular pyruvate is the key metabolite that triggers biofilm formation and identify a *C. difficile* pyruvate importer involved in this process.

## Results

### Overview of the time course transcriptomic analysis of *C. difficile* grown in the presence of DOC

In our previous study, we observed that when grown in the presence of sub-inhibitory concentrations of DOC, *C. difficile* enters stationary phase between 14 h and 20 h and by 48h has formed a strong biofilm (Dubois et al 2019). To identify key events leading to biofilm formation in the presence of DOC, we performed a time course transcriptomic analysis on *C. difficile* grown in BHI supplemented with yeast extract, cysteine, glucose (BHISG) and 240 µM DOC. For this analysis, time points were selected based on the growth curve. Planktonic bacterial cells were harvested in logarithmic phase (9 h), transition phase/early stationary phase (14 h), and mid-stationary phase (24 h); biofilm cells were harvested at 48 h. We then compared the expression profile of the cells at 14 h, 24 h and 48 h using the 9 h time point as our refence point. Relative to the 9 h time point, a total of 262, 659 and 659 genes were down-regulated at 14 h, 24 h and 48 h, respectively, whereas a total of 218, 794 and 997 genes were up-regulated at 14h, 24h and 48h, respectively (Supplementary data). We started our analysis by focusing on cell surface structures known to contribute to biofilm formation. The genes encoding type IVa pili (T4aP) machinery [*pilA2* (*CD630_32940*), *pilW* (*CD630_23050*) and the *pilA1* cluster (*CD630_35030-CD630_35130*)] were upregulated in the DOC-induced biofilm cells (Supplementary data).

### Cell surface proteins and structures play a limited role in biofilm formation

To see if the T4aP was critical for biofilm formation in the presence of DOC, the *pilA*2 cluster (*CD630_32910-CD630_32970*), the major pilin *pilW* and the *pilA1* cluster were deleted. Deletion of the *pilA1* cluster resulted in a significant decrease in biofilm formation but deletion of *pilW* or the *pilA2* cluster had no effect (Figure 1A). Given that the *pilA1* cluster was required for optimal biofilm formation, we tested the ability of a strain lacking the PilA1 major pilin to form biofilms (Poquet et al., 2018). Inactivation of *pilA1* (*CD630_35130*) did not affect biofilm formation (Supplementary Figure S1A).

**Figure 1:**
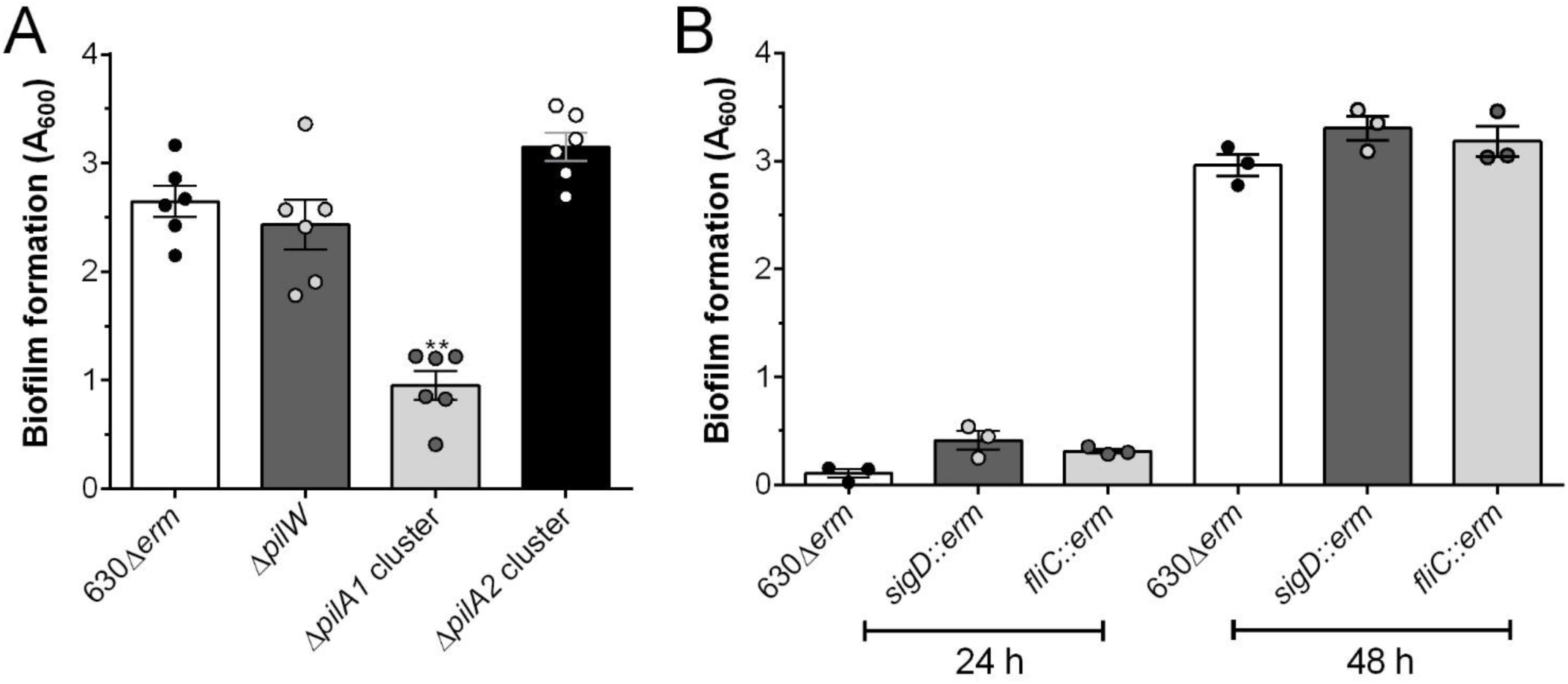
The T4aP machinery is required for biofilm formation in BHISG supplement with DOC. Biofilm formation was assessed in strains lacking genes (A) encoding pili (48h), (B) or involved in flagella motility (24h and 48h). Asterisks indicate statistical significance determined with a Kruskal-Wallis test followed by an uncorrected Dunn’s test (** *p*≤0.01, vs 630Δ*erm*). Data shown indicate biological replicates from experiment performed on different days, the bars represent the mean and the error bars represent the SEM.

T4aP gene expression is also under the control of c-di-GMP and overproduction of c-di-GMP results in measurable auto-aggregation and biofilm formation (Soutourina et al., 2013). Several c-di-GMP associated genes were up-regulated in biofilms (48 h) suggesting this signalling molecule might have a role (Supplementary Table 1 and Supplementary data). We then decided to test the effect of c-di-GMP overproduction on auto-aggregation in the presence of DOC. To do so, a *C. difficile* strain (CDIP634) with an inducible diguanylate cyclase (*CD630_1420*) that overproduces c-di-GMP was grown in the presence or absence of DOC. When we compared CDIP634 to a parental control strain, we observed high levels of auto-aggregation in BHISG exclusively when c-di-GMP was overproduced (Supplementary Figure S1B). However, auto-aggregation was not observed in BHISG supplemented with DOC, suggesting that cells are insensitive to c-di-GMP overproduction under our biofilm forming conditions (Supplementary Figure S1B). In addition to the T4aP, several genes associated with cell surface proteins and structures were differentially regulated in our transcriptome analysis (Supplementary data). Among those, the collagen binding protein gene (*CD630_28310*) was up-regulated at 24 h and 48 h, and *bcsA* (*CD630_25450*), which encodes a protein with a cellulose synthase domain, was up-regulated at 48 h (Supplementary data). Given that CD630_28310 acts as an adhesin and BcsA could promote exopolysaccharide synthesis, we tested the ability of strains lacking either gene to form biofilm in the presence of DOC. Absence of either gene did not affect biofilm formation (Supplementary Figure S1A).

In our transcriptome data, we also observe that the F1 (late), F2 (glycolysation) and F3 (early) clusters of the flagellum biogenesis gene clusters were down-regulated at 48 h (Supplementary data). Therefore, we tested the biofilm forming kinetics of a strain lacking the flagellin *fliC* and a strain lacking the alternative sigma factor for the flagellar operon *sigD* to see if inactivation of flagellum synthesis affects biofilm formation.

Inactivation of *fliC* or *sigD* had no effect on biofilm formation or its kinetics (Figure 1B). This suggests that the flagellum is not required for biofilm formation and that absence of the flagellum does not enhance biofilm formation.

Altogether, we show that the T4aP machinery encoded by the *pilA1* cluster is required for biofilm formation in the presence of DOC. Other surface proteins associated with biofilm formation in the absence of DOC (Dapa et al., 2013; Poquet et al., 2018) are dispensable, suggesting a distinct biofilm-formation mechanism.

### Biofilm formation in the presence of DOC is associated with profound metabolic rearrangement

We then analysed our time-course transcriptomic data using the Biocyc database for changes in metabolic pathways (see Supplementary Figure S2). As observed in our previous study (Dubois et al. 2019), genes encoding enzymes associated with glycolysis were down-regulated from 14 h onward (see Supplementary Figure S2 and Supplementary data). Furthermore, enzymes involved in reductive and oxidative Stickland reactions were up-regulated at 14 h and 24 h, but their expression was unchanged at 48 h. Genes associated with fermentation pathways such as butanoate were up-regulated at 24h but down-regulated at 48h (see Supplementary Figure S2 and Supplementary data). Interestingly, *C. difficile* appears to down-regulate genes associated with ethanolamine degradation (see Supplementary Figure S2 and Supplementary data), a valuable nutrient source that is converted to acetyl-CoA (Nawrocki et al., 2018).

We also observed that PTS dependent glucose transporters were down-regulated from 14 h onwards but a predicted glucose transporter (CD630_30170) was up-regulated at 48 h. Different PTS-dependent transporters predicted to import sugars other than glucose were up-regulated at different time point. Specifically, the mannitol importer genes (MtlFA) and the gene encoding a mannitol-1-phosphate 5-dehydrogenase (MtlD) were up-regulated at every time points (see Supplementary Figure S2 and Supplementary data). The PTSs predicted to transport fructose (CD630_30130-CD630_3015 and CD630_24860-CD630_2488) and/or mannose (CD630_30670-CD630_3069) were up-regulated at 48 h but the PTS-dependent fructose transporter FruABC was down-regulated at 14h and 24h (see Supplementary Figure S2 and Supplementary data). However, genes encoding enzymes processing fructose (FruK) or mannose (Pmi and CD630_23180) were down-regulated at 14 h and 24h or 48h, respectively (see Supplementary Figure S2 and Supplementary data). Gene encoding PTS-dependent transported predicted to import maltose or N-acetylglucosamine were up-regulated at different time points. For example, CD630_04690 was only up-regulated at 14h but CD630_13360 was up-regulated at 48 h. However, genes encoding enzymes processing maltose or GlcNAc were not differently regulated (see Supplementary Figure S2 and Supplementary data). Furthermore, the genes encoding proteins that import and process N-acetylneuraminic acid (Neu5Ac) were up-regulated at 48 h (see Supplementary Figure S2 and Supplementary data). This could indicate that *C. difficile* might use different carbon source once glucose is depleted, probably near the 14h time point.

We then compared our transcriptome to a published omics analysis of *C. difficile* cells in different growth phases (Hofmann et al., 2018). Based on these comparisons, we found evidence that the cells harvested at 24 h have a transcriptome profile of stationary phase cell, (Figure 2). Specifically, transcription of genes associated with protein degradation, butanoate fermentation, acetate fermentation, glycine metabolism, and oxidative Stickland reaction of branched chained amino acids were up-regulated at 24 h (Figure 2 and Supplementary data) as observed for cells in the stationary phase (Hofmann et al., 2018). Expression of these genes decreased at 48 h (Figure 2). Other genes that were up-regulated during stationary phase were those associated with cysteine biosynthesis, pantothenate biosynthesis, riboflavin biosynthesis, ferrous iron transport, flavodoxin and chaperones (Figure 2) but these genes remain induced at 48 h in our analysis (Supplementary data). On the other hand, genes associated with glycolysis were down-regulated at 24 h as observed for the stationary phase analysis and their expression remains down-regulated in biofilm cells at 48 h (Figure 2). Interestingly, genes involved in protein synthesis were down-regulated at 24 h but their expression increased at 48 h (Figure 2 and Supplementary data).

**Figure 2:**
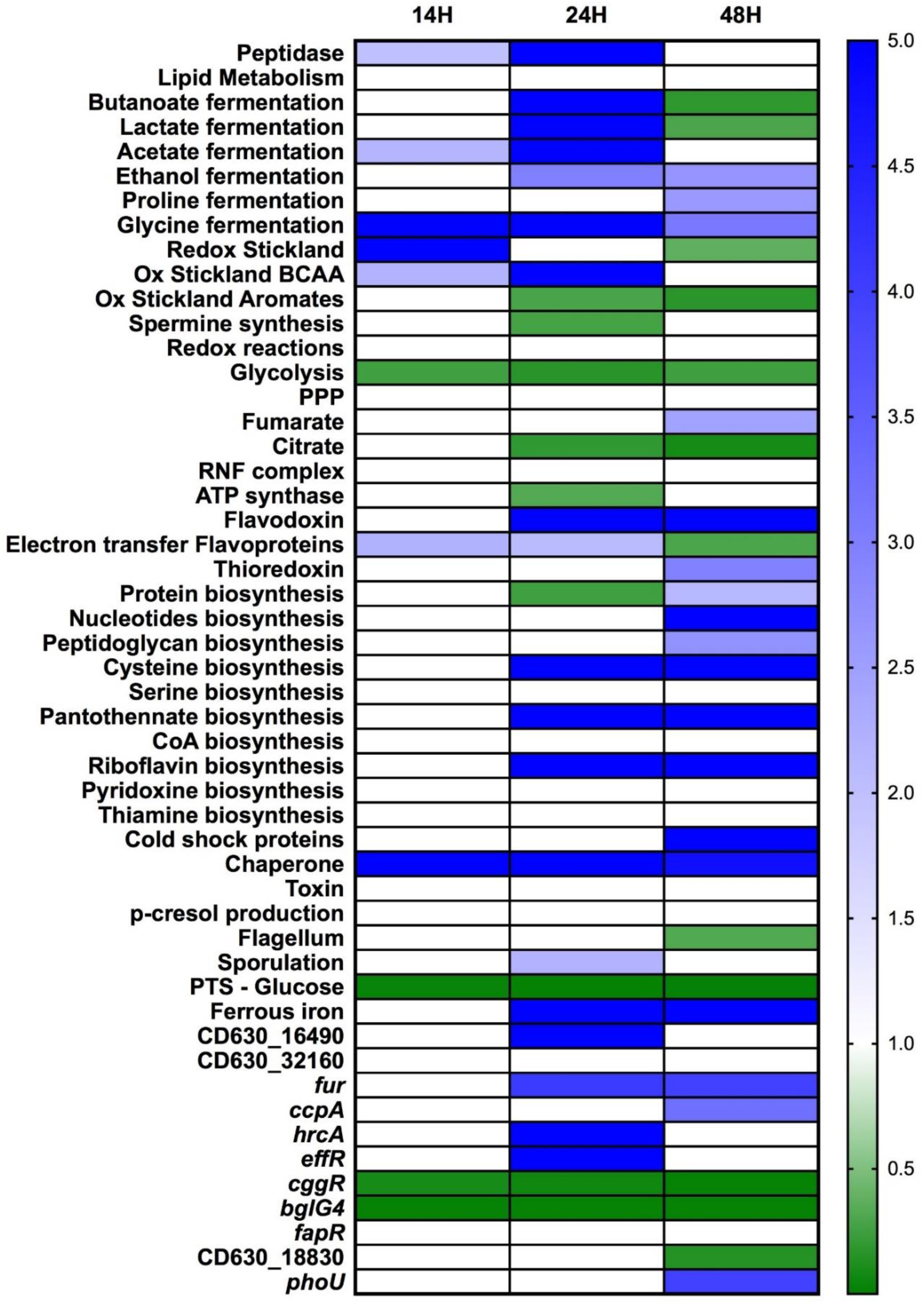
Overview of transcriptomic changes over time in genes previously associated with stationary phase in *C. difficile* strain 630Δ*erm* grown in BHISG supplemented with 240 µM DOC. Changes in expression over time are colour coded (in white, no changes; blue, up-regulated; or green, down-regulated). The 9 h time point was used as the reference point to measure gene expression. Data used to generate the figure are available in the Supplementary data file

Overall, these metabolic changes overlap with our previous analysis comparing cells grown in BHISG in the presence or absence of DOC at 48h (Dubois et al., 2019). For example, we observe at 48h down-regulation of genes involved in glycolysis and up-regulation of genes predicted to encode transporters of alternate carbon source (see Supplementary Figure S2 and Supplementary data). This clearly supports the idea that DOC induces a metabolic stress in *C. difficile* and long-term exposure to DOC results in a remodelling of the metabolic profile of *C. difficile*. We also clearly see that the 24 h transcriptomic signature of *C. difficile* grown in BHISG in the presence of DOC resembles cells in a stationary phase but at 48 h, once *C. difficile* is in a biofilm state, metabolic activity and transcriptional activity increase.

### The transition phase regulator SigH is required for DOC-induced biofilm formation

In *C. difficile*, SigH and SinR modulate Spo0A expression and activity in addition to mediating the transition from exponential to stationary phase (Saujet et al., 2011; Girinathan et al., 2018). Therefore, based on the observation that *spo0A* inactivation decreases biofilm formation in the presence of DOC (Dubois et al., 2019; Figure 3A) in combination with our time course transcriptomic analysis indicating the effect of major transcriptional changes associated with stationary phase, we tested the effect of SigH and SinR on biofilm formation. Inactivation of *sigH* results in decreased biofilm-formation whereas inactivation of *sinR* did not have an effect (Figure 3A and Supplementary Figure S3A).

**Figure 3:**
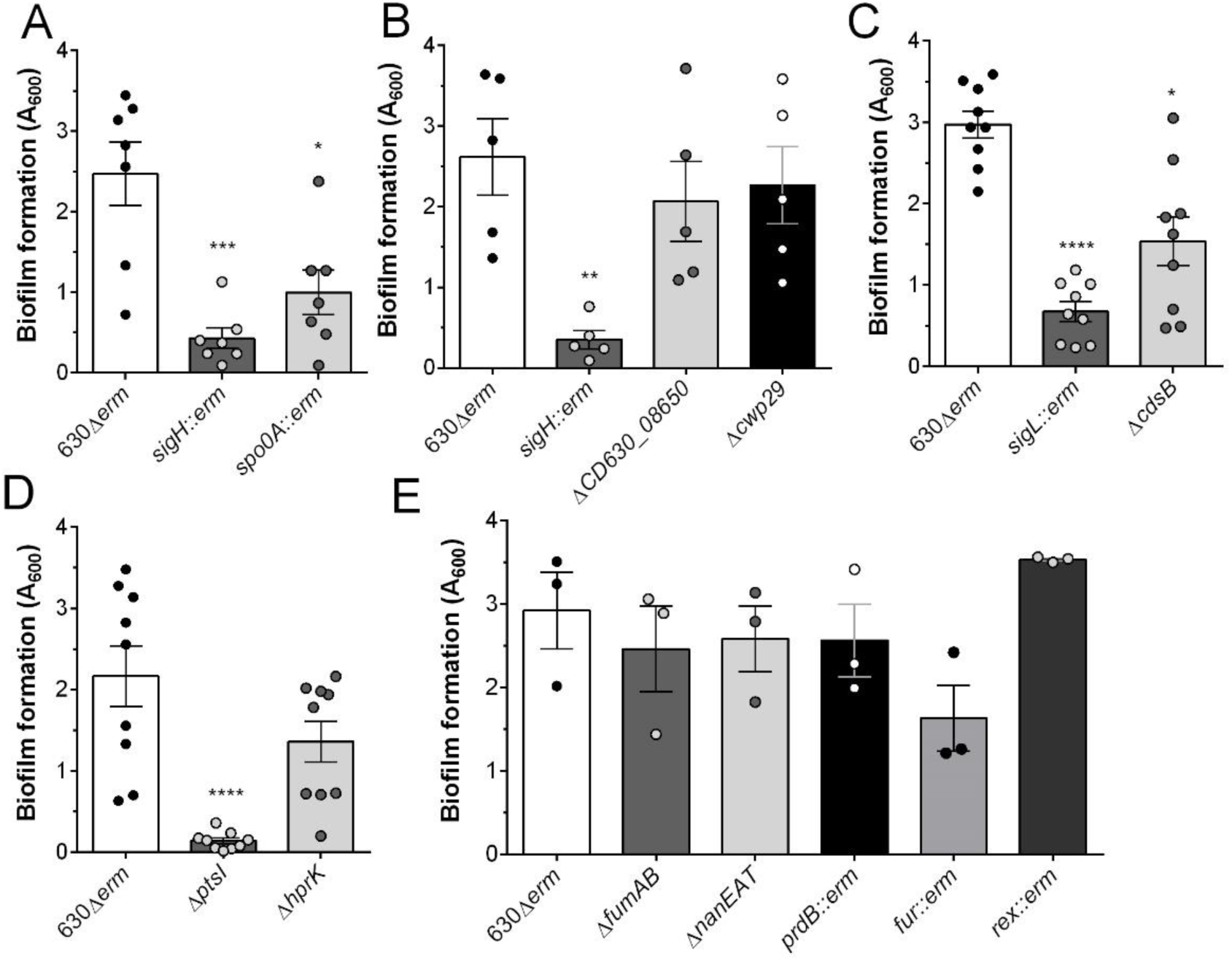
Biofilm formation in BHISG supplement with DOC is dependent on SigH, SigL and PTS transport. Biofilm formation was assessed 48h after inoculation in strains lacking genes involved in (A) the transition to stationary phase, (B) the *sigH* regulon, (C) cysteine metabolism, (D) PTS transport and (E) other metabolic pathways. Asterisks indicate statistical significance determined with a Kruskal-Wallis test followed by an uncorrected Dunn’s test or a two way ANOVA followed by a Fisher’s least significant difference test (** *p*≤0.001 *** *p*≤0.001, **** *p*≤0.0001vs 630Δ*erm*). Data shown indicate biological replicates from experiment performed on different days, the bars represent the mean and the error bars represent the SEM.

Inactivation of *sigH* decreased biofilm formation to a lower level than observed in the *spo0A* inactivated strain. Given that our time-course transcriptome did not show changes in *spo0A* expression, our data suggest that genes other than *spo0A* that are under SigH control contribute to biofilm formation. To identify SigH controlled genes that contribute to biofilm formation, we compared the list of genes with a SigH binding site in their promoter (Saujet et al., 2011) with our transcriptome and identified specific genes that were up-regulated at 24 h (Supplementary Table 2). We selected 4 genes (*CD630_08650*, *CD630_12640*, *cwp29* and *CD630_34580*) that were not involved in sporulation, transcription or metabolism and we tested their expression in the *sigH*::*erm* strain and the parental strain grown in BHISG with DOC at 24 h. The expression of *CD630_08650* and *cwp29* was greatly reduced in the absence of SigH whereas the expression of *CD630_12640* and *CD630_34580* was only moderately reduced (Supplementary Figure S3B). This confirmed that expression of *CD630_08650* and *cwp29* is controlled by SigH in our biofilm inducing condition.

Based on these results, we deleted *CD630_08650* and *cwp29* and tested the ability of the resulting strains to form biofilm. Deletion of either gene did not have an effect on biofilm formation (Figure 3B). Unfortunately, efforts to generate a strain lacking both *CD630_08650* and *cwp29* were unsuccessful. When we tested the viably of the *sigH*::erm strain, this strain had a one log reduction in viability compared to the parental strain (Supplementary Figure S3C). The viability of the *spo0A*::*erm* strain is not affected in our conditions (Dubois et al., 2019). The decrease in viability of the *sigH*::*erm* strain provides a possible explanation for the reduction in biofilm formation. Based on our previous study, we know that biofilm formation in the presence of DOC is not dependent on sporulation (Dubois et al., 2019). In addition, SigH and Spo0A contribute to metabolic adaptation of *C. difficile* (Saujet et al., 2011; Pettit et al. 2014). Taken together, the sub-optimal metabolic profile of these strains might result in a decrease in biofilm formation highlighting the impact of metabolism in *C. difficile* persistence.

### PTS mediated transport and cysteine metabolism are required for DOC-induced biofilm formation

Given our transcriptomic data shows changes in metabolism, we next tested the effect of inactivating different metabolism-associated genes. We first tested the role of cysteine metabolism because several genes associated with this metabolic pathway were differentially regulated in our transcriptomic analysis (Figure 2 and Supplementary data). Inactivation of the OAS-thiol-lyase encoding gene *cysK*, which participate in cysteine biosynthesis, did not alter biofilm formation (Supplementary Figure S4A). We then tested inactivation of the sigma factor encoding gene, *sigL* and deletion of the SigL-controlled cysteine desulfidase encoding gene, *cdsB*, important for amino acid degradation and cysteine catabolism resulting in the production of pyruvate and sulfide, respectively (Dubois et al., 2016 and Gu et al., 2018). Inactivation of *sigL* greatly reduced biofilm formation by *C. difficile,* whereas deletion of *cdsB* had an intermediate phenotype that was highly variable (Figure 3C). Complementation of the *sigL*::erm and Δ*cdsB* strain with *sigL* or *cdsB* expressed from their native promoter on a plasmid restored biofilm formation (Supplementary Figure S4B).

We then targeted the phosphotransferase system (PTS) and carbon catabolite repression (CCR) regulatory network by deleting *ptsI* that encodes enzyme I of the PTS and deleting *hprK* that encodes the Hpr kinase which is involved in carbon metabolism regulation and transport. Biofilm formation was abolished in the Δ*ptsI* strain whereas only a slight reduction in biofilm formation was observed in the Δ*hprK* strain (Figure 3D). Complementation of the Δ*ptsI* strain with inducible *ptsI* on a plasmid restored biofilm formation (Supplementary Figure S4B). These observations support our previously published results (Dubois et al., 2019) showing that the presence of glucose and carbon metabolism are essential for DOC-induced biofilm formation. However, there might be some redundancy within the carbon metabolism regulation network because deletion of *hprK* had limited and variable effects. These findings are in line with our transcriptomic analysis indicating that a profound metabolic reorganisation occurs prior to DOC-induced biofilm formation. Given that biofilm formation and glucose transport are both PTS dependent, our data suggest that *C. difficile* must first use glucose and then switch its metabolic profile to use different metabolic pathways, including those dependent on SigL, SigH and Spo0A to induce biofilm formation in the presence of DOC.

To elucidate the downstream metabolic pathways driving biofilm-formation, we targeted genes identified in our transcriptomic analysis associated with the conversion of fumarate to pyruvate (*fumAB-CD630_10500*), Neu5Ac transport and metabolism (*nanEAT*), proline metabolism (*prdB*), low intracellular NADH/NAD+ activated regulator (*rex*) and the ferric uptake regulator (*fur*). We then tested the ability of these gene deletion and inactivation strains to form biofilms. Every strain tested formed biofilms similar to the parental strain (Figure 3E) with the exception of the *fur*::*erm* strain that had a slight growth delay in BHISG+DOC while biofilm levels reached that of the parental strain by 72 h (data not shown). Taken together, our results suggest that BHISG provides multiple and redundant nutrient sources and only specific nutrients such as glucose and cysteine are essential for DOC-induced biofilm formation in *C. difficile*.

### Branched-chain amino acids and mucus-derived sugars potentiate the effect of DOC

The use of a complex medium (BHISG) in the biofilm formation assay made it difficult to identify specific metabolic processes and metabolites involved in DOC-induced biofilm formation. Therefore, we sought to optimize a minimal medium that could support biofilm formation in the presence of DOC. We first tested the *C. difficile* minimal medium (CDMM) described by Cartman and Minton (2010) with some modifications. Specifically, glucose and cysteine were increased to 100 mM and 0.1%, respectively, to match the concentrations present in our complex medium. We also tested different sources of amino acids and/or peptides. To increase the biofilm biomass, we used our transcriptomic data to identify key metabolic pathways that were up-regulated during biofilm formation and observed that predicted branched chain amino acid (BCAA) transporters (*CD630_12590*, *CD630_12600*, *CD630_27020*) were up-regulated at 24 h.

Therefore, we added BCAA to our semi-defined medium and observed an increase in biofilm biomass when the medium was supplemented with BCAA, cysteine and a carbohydrate source, such as glucose (Figure 4A, 4B and Supplementary Figure S5A). We noted that only casein hydrolysate from Oxoid supported biofilm formation while casamino acids from Difco or a mixture of individual essential amino acids did not (Supplementary Figure S5A). The resulting medium that supported biofilm formation was named *C. difficile* Medium Optimized for Biofilm formation (CDMOB). In addition to BCAA transporters being up-regulated during biofilm formation, the Neu5Ac transporter was also up-regulated in our transcriptomic analysis (Supplementary data). Thus, we tested if mucus-derived sugars potentiate DOC-induced biofilm formation. Mucus is typically broken down into different hexose sugars including glucose, GlcNAc, fucose, Neu5Ac, galactose and N-acetylgalactosamine (GalNAc) and GlcNac and Neu5Ac acquisition is important for *C. difficile* growth in the intestinal tract (Ng et al., 2013; Pereira et al., 2020). We tested the effect of these sugars on biofilm formation. When CDMOB was supplemented with DOC and 100 mM of each sugar; the addition of glucose, GlcNAc or Neu5Ac induced biofilm formation, whereas addition of fucose, galactose and GalNAc had no effect (Figure 4C). We then sought to see if combining different sugars has additive effects on biofilm formation. When glucose and GlcNAc were mixed at concentrations below those required for biofilm formation (100 mM), the mixture supported biofilm formation at levels equivalent to those with a single sugar (Figure 4D). We then tested the biofilm-formation ability of a strain lacking the *nanEAT* operon in the presence of 100 mM Neu5Ac. This operon encodes a non-PTS transporter (*nanT*), an acetylneuraminate lyase (*nanA*) and the N-acetylmannosamine-6-phosphate 2-epimerase (*nanE*), involved in the metabolism of Neu5Ac (Supplementary Figure S2). In the absence of *nanEAT* but not *ptsI*, *C. difficile* failed to form a biofilm in CDMOB supplemented with DOC and 100 mM Neu5Ac (Figure 4E). Taken together, our results reinforce the idea that metabolized sugars (glucose, GlcNAc and Neu5Ac) potentiate the effect of DOC (see Dubois et al 2019).

**Figure 4:**
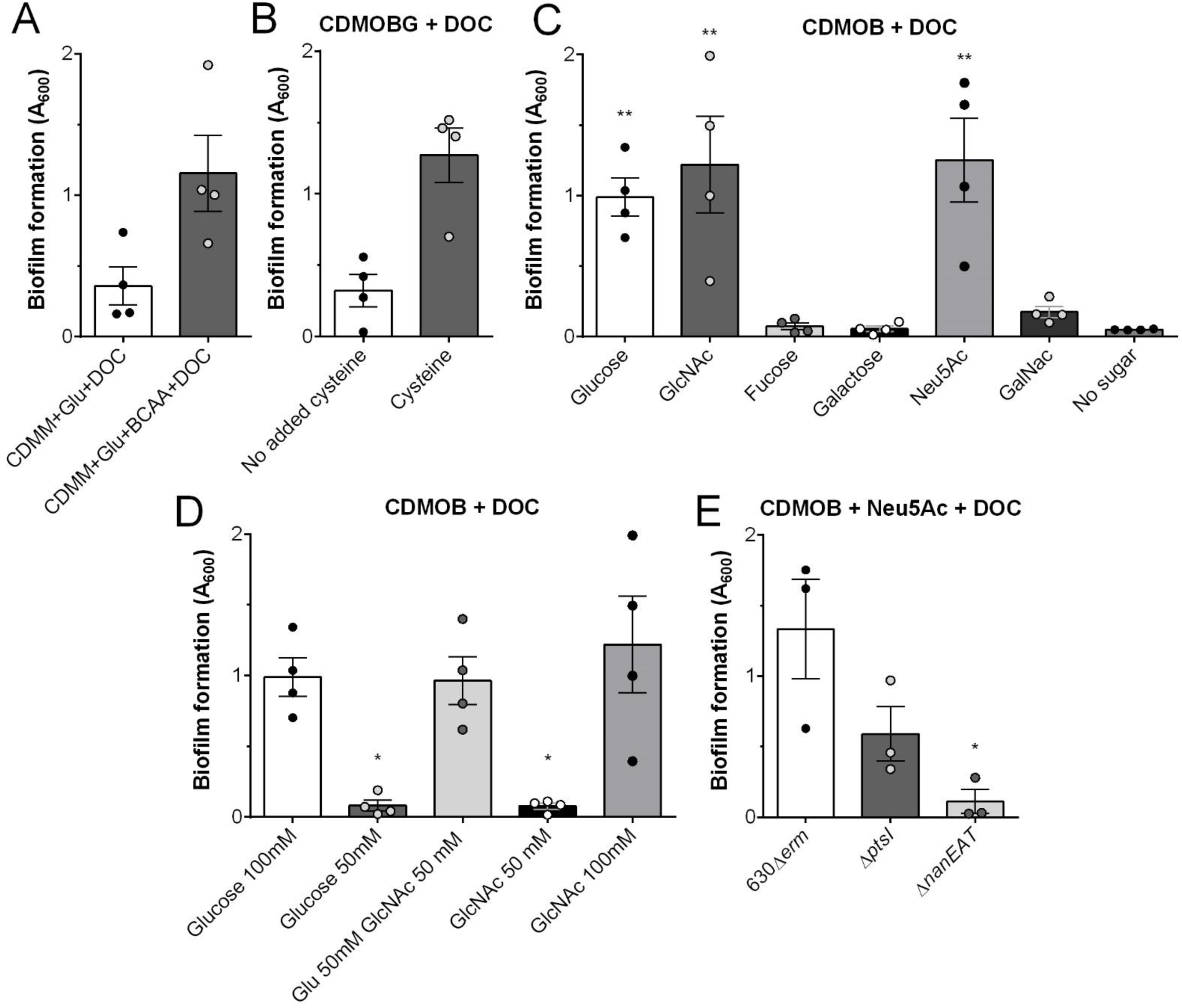
Combination of branched chain amino acid, cysteine and mucus derived sugars induce biofilm formation in CDMOB supplemented with 240 µM DOC. Biofilm formation was assessed 48 h after incubation when medium was supplemented with 1% (w/v) BCAA supplementation (A), without cysteine supplementation (B),100 mM mucus derived sugars (C) or glucose (Glu) and N-acetyl-glucosamine (GluNAc; D). Neu5Ac: N-acetylneuraminic acid; GalNAc; N-acetylgalactosamine. Experiments presented in panel B and C were performed at the same time. Biofilm formation by Δ*ptsI* and Δ*nanEAT* in medium supplemented with 100 mMNeu5Ac (E). Asterisks indicate statistical significance determined with a Kruskal-Wallis test followed by an uncorrected Dunn’s test (* *p*≤0.05, ** *p*≤0.001 vs no sugars, 100 mM glucose or 630Δ*erm*). Data shown indicate biological replicates from experiment performed on different days, the bars represent the mean and the error bars represent the SEM.

To further characterize the biofilm formed in CDMOB supplemented with glucose (CDMOBG) and DOC, we analysed the composition of the biofilm matrix by enzymatic dispersion of preformed biofilms and by gel electrophoresis of the isolated matrix. As observed previously with biofilm formed in BHISG supplemented with DOC (Dubois et al 2019), NaIPO_4_ treatment, which denatures polysaccharides, failed to disperse preformed biofilms, while DNase-treatment dispersed preformed biofilms (Supplementary Figure S5B). Unlike what was observed in BHISG with DOC, proteinase K treatment dispersed preformed biofilms from cells grown in CDMOBG with DOC. When we analysed biofilm matrix of cells grown in CDMOBG with DOC, we observed extracellular DNA (eDNA) and patterns of proteins, glycoproteins and DNase/proteinase resistant material (e.g. polysaccharides or glycosylated amyloid-like fibers) that was similar to that of the biofilm matrix of *C. difficile* grown in BHISG with DOC (Supplementary Figure S5C, D and E). However, the high molecular weight smear observed in BHISG with DOC disappeared from the matrix of biofilms from cells grown in CDMOBG with DOC (Supplementary Figure S5E and S5F). This disappearance suggests that either *C. difficile* produces high molecular weight glycol molecules in BHISG with DOC or that the smear is from component(s) of the medium in our samples. Taken together, our data indicate that the biofilm formed in CDMOBG with DOC is similar to the one formed in BHISG with DOC but proteins play a larger role in maintaining biofilm matrix stability when *C. difficile* is grown in CDMOBG with DOC.

### Extracellular pyruvate is required for biofilm formation in CDMMBG with DOC

Given that we detect biofilm formation at 48h but not 24h, we hypothesized that *C. difficile* detects high-cell density via quorum sensing. In support of this hypothesis, we observed up-regulation at 24 h of the genes encoding the autoinducer *agr* and its associated transporter protein, while *luxS* was down-regulated (Supplementary data). Deletion of the autoinducer *agr* operon or inactivation of *luxS* did not alter DOC-induced biofilm formation (Supplementary Figure S4A).

Since deletion of typical quorum sensing molecules did not appears to alter DOC-induced biofilm formation, we hypothesized that a metabolite could drive this lifestyle switch. We analysed the volatile and non-volatile acid content of spent culture supernatants. After 24 h of growth in BHISG with DOC, pyruvic acid levels were high and these levels were decreased at 48 h. In the absence of DOC, pyruvic acid levels were lower, but the level did not decrease over time (Supplementary Figure S6A).

Based on this analysis, we hypothesized that high level of extracellular pyruvate was important for initiating DOC-induced biofilm formation. To build on this observation, we tested the effect of pyruvate depletion on biofilm formation by adding pyruvate dehydrogenase to *C. difficile* cultures. When extracellular pyruvate was enzymatically depleted by pyruvate dehydrogenase addition at 24 h in CDMOBG with DOC, biofilm formation was inhibited. This effect was not observed when heat inactivated pyruvate dehydrogenase or buffer were added to CDMOBG with DOC at 24h (Figure 5A). Based on these data, we conclude that pyruvate might act as a critical molecule triggering *C. difficile’s* switch from a planktonic to a biofilm lifestyle in the presence of DOC.

**Figure 5:**
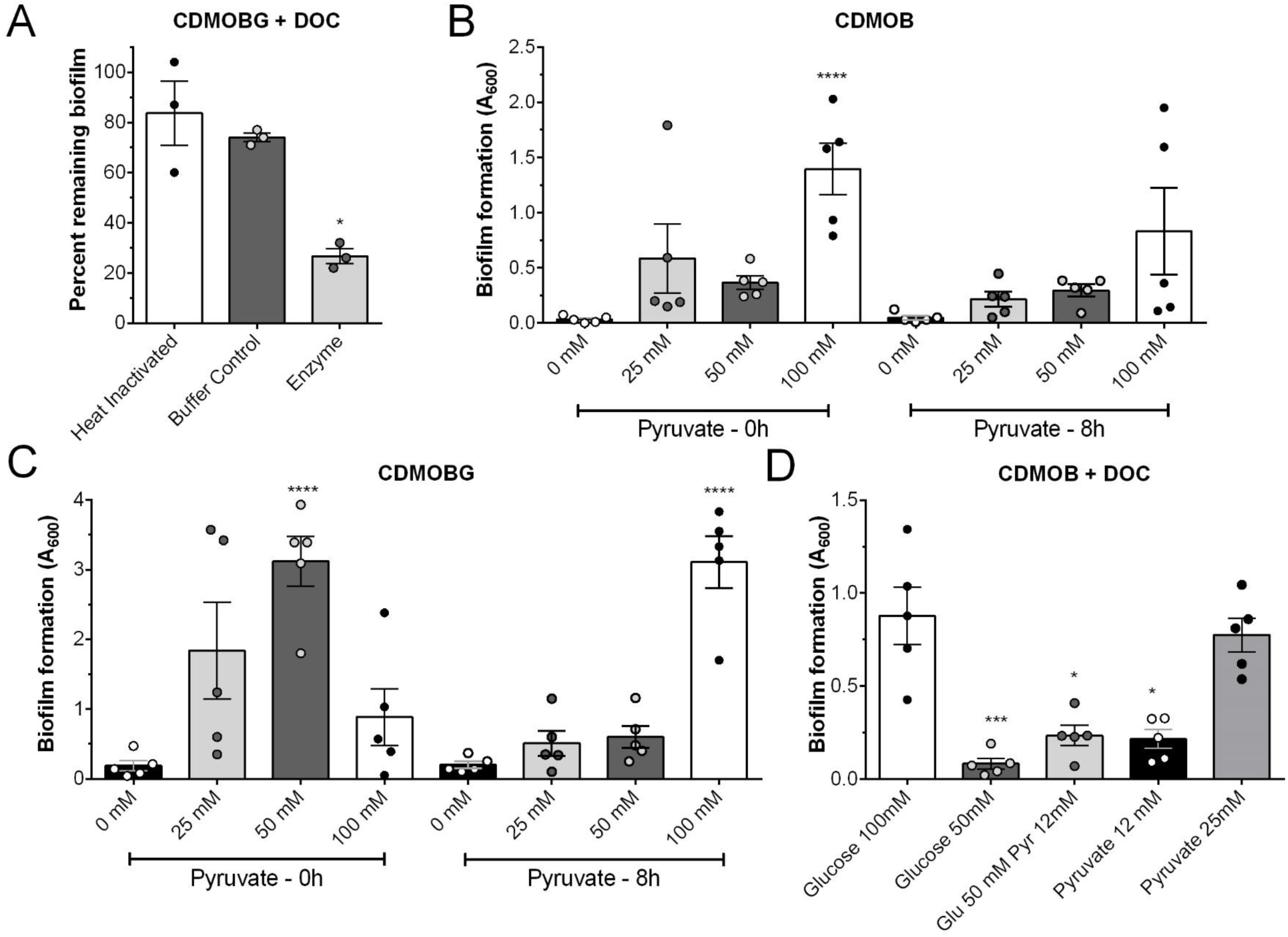
Presence of extracellular pyruvate induces biofilm formation. (A) Effect of enzymatic depletion of extracellular pyruvate on 48 h biofilm formation in *C. difficile*. Pyruvate dehydrogenase was added to *C. difficile* grown in CDMOBG after 24 h of growth. Control culture were treated with heat inactivated enzyme or buffer are shown. Effect of pyruvate on 48 h biofilm formation in the absence of DOC by *C. difficile* grown in CDMOB (B) or CDMOBG (C). Where indicated, pyruvate was added at inoculation (0 h) or after 8 hours of growth (8 h). (D) Biofilm formation for *C. difficile* 630Δ*erm* in CDMOB in the presence of 240 µM DOC and glucose (glu) and/or pyruvate (pyr). For A, and D, asterisks indicate statistical significance determined with a Kruskal-Wallis test followed by an uncorrected Dunn’s test (* *p*≤0.05, vs heat inactivated enzyme; * p≤0.05, *** p≤0.001 vs 100 mM glucose). For B, and C, asterisks indicate statistical significance determined with a two-way ANOVA followed by Fisher’s least significant difference test (**** *p*≤0.001 vs 0 mM pyruvate). Data shown indicate biological replicates from experiment performed on different days, the bars indicate the mean and the error bars represent the SEM.

### Pyruvate induces biofilm formation in the absence of DOC

To test if pyruvate alone could induce biofilm formation, we supplemented CDMOB or CDMOBG with increasing concentrations of pyruvate (25, 50 and 100 mM). When added at inoculation, pyruvate induced biofilm formation in CDMOB and CDMOBG at 100 mM and 50 mM respectively (Figure 5B and 5C). When it was added after 8h of growth (log to stationary phase transition), pyruvate only induced biofilm formation in CDMOBG at 100 mM (Figure 5C). We also added pyruvate after 24 h of growth but this failed to induce biofilm formation (Data not shown). This indicates the metabolic state of the bacteria partially determine whether pyruvate will induce biofilm formation. We then tested if pyruvate could cooperate with DOC to support biofilm formation and observed that 25 mM, instead of 100 mM pyruvate was sufficient to support biofilm formation in CDMOB with DOC (Figure 5D). However, glucose and pyruvate did not have a cooperative effect when added at 50 mM and 12.5 mM, respectively (Figure 5D). When we consider our biofilm formation assay, the gas chromatography data and our pyruvate depletion assay, our results indicate that biofilm-formation depends on the amount of available pyruvate and suggests that this metabolite is a key factor driving DOC-induced biofilm formation.

### Induction of biofilm formation requires pyruvate sensing by the *CD630_26010-26020* TCS and pyruvate importer CstA

In a previous study, we identified a LytRS two-component regulatory system (TCS) homologue (CD630_26020-CD630_26010) that regulated toxin gene expression in response to pyruvate (Dubois et al. 2016). In other Gram-positive bacteria, genes encoding TCSs sensing pyruvate are associated with a pyruvate importer and a gene encoding a potential importer (*CD630_26000*) that is located immediately downstream of *CD630_26010*. CD630_26000 encodes a CstA homologue that was initially annotated as a peptide transporter, but homologues were recently shown to be pyruvate importers in *Escherichia coli* (Hwang et al. 2018). To determine if CstA or TCS CD630_26020-26010 are involved in biofilm formation and respond to pyruvate availability, we deleted *cstA* and tested whether the Δ*cstA* strain and the previously described CD630_26020::*erm* strain (Dubois et al., 2016) form biofilms in CDMOB with 100 mM pyruvate and CDMOBG with 50 mM pyruvate. Pyruvate was added at inoculation and the Δ*ptsI* strain, unable to uptake PTS-dependent sugars, was used as a positive or negative control when pyruvate was added alone or with glucose, respectively, to the growth medium.

In CDMOB supplemented with 100 mM pyruvate as the sole carbon source, the *CD630_26020*::*erm* and Δ*cstA* strains did not form biofilm whereas, as expected, the Δ*ptsI* and parental strain formed biofilms (Figure 6A). In CDMOBG supplemented with 50 mM pyruvate, the *CD630_26020*::*erm* and Δ*cstA* strains formed biofilms whereas the Δ*ptsI* strain did not form a biofilm (Figure 6B). The inability of the Δ*ptsI* strain to form a biofilm in the presence of glucose confirms that glucose metabolism is required for biofilm formation. This might be due to the production of pyruvate by glycolysis which would increase the level of extracellular pyruvate above the threshold required to induce biofilm-formation (Figure 5A and C). Absence of biofilm formation for the Δ*cstA* strain or the *CD630_26020*::*erm* strain when pyruvate is the sole carbon source suggests that the CstA importer and the CD630_26020-CD630_26010 TCS are primarily involved in the uptake and sensing of extracellular pyruvate, respectively. Biofilm formation by the Δ*ptsI* strain indicates that the PTS system is not involved in pyruvate uptake.

**Figure 6:**
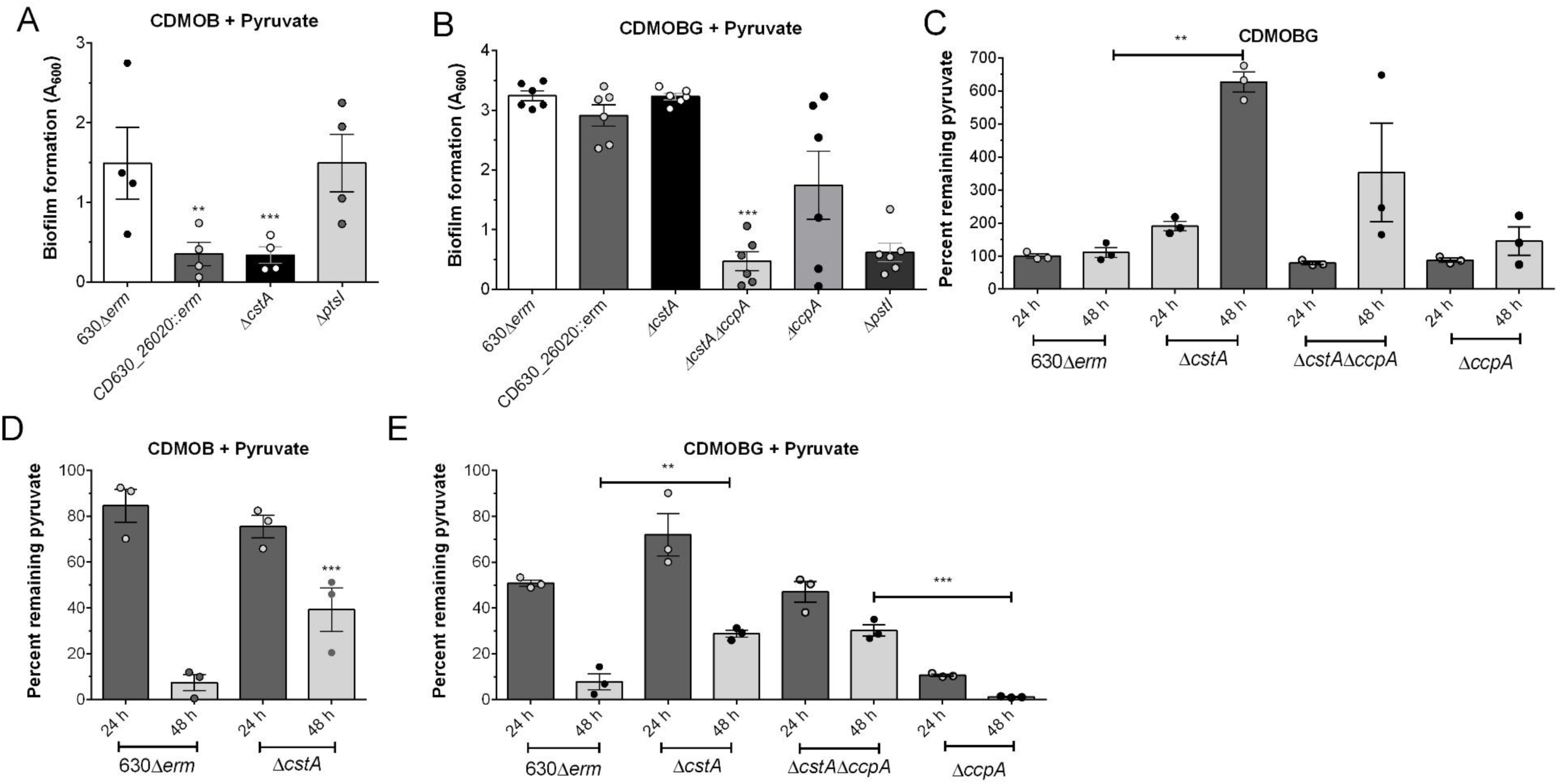
Biofilm formation in the presence of pyruvate is CtsA dependent. Biofilm formation was assessed in strains lacking the CD630_26020-CD630_26010 TCS, *cstA*, *ptsI*, *ccpA* or *ccpA* and *cstA* in CDMOB with pyruvate (A) and CDMOBG with pyruvate (B). Pyruvate used during growth by the parental, Δ*cstA*, Δ*ccpA* or Δ*cstA*Δ*ccpA* strains grown in CDMOBG (D), CDMOB with 100 mM pyruvate (D) or CDMOG with 50 mM pyruvate (E) (% remaining = 100 × pyruvate in media after growth (24 h or 48 h)/ starting pyruvate concentration). For A, and B, asterisks indicate statistical significance determined with a Kruskal-Wallis test followed by an uncorrected Dunn’s test (** *p*≤0.001 *** *p*≤0.001, vs 630Δ*erm*). For C, D and E, asterisks indicate statistical significance determined with a two-way ANOVA test followed by Fisher’s least significant difference test (** *p*≤0.001, *** *p*≤0.001 vs 630Δ*erm*). Data shown indicate biological replicates from experiment performed on different days, the bars represent the mean and the error bars represent the SEM.

The ability of the Δ*cstA* strain or the *CD630_26020*::*erm* strain (Figure 6B) to form biofilms in the presence of glucose and pyruvate suggests that at least in the presence of glucose, CstA or CD630_26020-CD630_26010 are not solely required for the uptake of pyruvate and that other importer(s) and/or regulatory mechanisms might be involved. In other bacteria, multiple proteins have been identified as pyruvate importers and expression of these importers are often controlled by the CCR regulatory network (Charbonnier et al. 2017; van den Esker et al. 2017). In *C difficile*, CcpA is the major regulator of CCR, which controls the use of alternate carbon sources. In support of its role in controlling pyruvate metabolism, CcpA binds to the region upstream of *cstA* (Antunes et al. 2012). To test if the catabolic repression system was involved in pyruvate transport, we deleted *cstA* in our Δ*ccpA* strain. The resulting double deletion strain (Δ*ccpA*Δ*cstA*) was tested for its ability to form biofilms in CDMOBG supplemented with 50 mM pyruvate. The Δ*ccpA*Δ*cstA* strain did not form a biofilm whereas the Δ*ccpA* or Δ*cstA* strains formed biofilms, although the amount of biofilm formed by the Δ*ccpA* strain was more variable (Figure 6B). These data support that extracellular pyruvate could also be imported by a second importer controlled by CcpA in the presence of glucose.

To test if extracellular pyruvate was used by the parental and deletion strains, pyruvate levels in the culture supernatant was measured from cells grown in CDMOB with 100 mM pyruvate, CDMOBG or CDMOBG with 50 mM pyruvate for 24 h and 48 h. In CDMOBG, our assay did not detect significant changes in extracellular pyruvate concentration in the parental strains but the Δ*cstA* strain accumulated 6 times more pyruvate in its supernatant (Figure 6C). This accumulation suggests that the Δ*cstA* strain can excrete pyruvate in a CstA-independent manner. In CDMOB with 100 mM pyruvate, the Δ*cstA* strain used approximately 50% of the pyruvate present in the growth medium but did not deplete pyruvate to the same extent as the parental strain (10% of starting concentration; Figure 6D). In CDMOBG with 50 mM pyruvate, the Δ*cstA* and Δ*ccpA*Δ*cstA* strains did not reduce the amount pyruvate to the levels of the parental or the Δ*ccpA* strains after 48h (Figure 6E). However, the Δ*cstA* and the Δ*ccpA*Δ*cstA* strain were able to reduce the amount of pyruvate present in the growth medium. Interestingly, the Δ*ccpA* strain was able to use 90% of the pyruvate by 24h. The rapid depletion of pyruvate in the Δ*ccpA* strain compared to the parental strain and the reduction in the amount of extracellular pyruvate used by the Δ*ccpA*Δ*cstA* strain suggest that CstA, and not another importer, is highly active in the absence of CcpA. This is consistent with previous data showing that CcpA binds to the region upstream of *cstA* and supports the hypothesis that *cstA* expression is repressed by CcpA in the presence of glucose (Antunes et al, 2012).

The decrease in extracellular pyruvate in the absence of CstA suggests that pyruvate is also imported independently of CstA (Figure 6E). In other bacteria such as *Bacillus subtilis,* there are additional pyruvate importers encoded by *pftAB* that belong to the LrgAB holin anti-holin family. When these predicted holin anti-holin systems act as pyruvate importers, their coding genes are invariably located next to a TCS. While an *lrgAB*-family gene (∼35% amino acid identity to PftB and no PftA homologs) exists in *C. difficile* strain 630, the genes are not associated with a TCS and were down-regulated at 24 h in BHISG supplemented with DOC (Supplementary data), suggesting that this *lgrB* gene homolog is unlikely to encode the additional pyruvate importer in *C. difficile*.

In addition to the *lrgAB* homolog, we noted the presence of a second *cstA-*homolog encoded by *CD630_23730*. This gene was highly expressed in *C. difficile* at 24 h in BHISG supplemented with DOC (Supplementary data); however, unlike *cstA* it is not located next to a TCS. It is possible that *CD630_23730* is the importer responsible for the partial pyruvate uptake by the Δ*cstA* and Δ*ccpA*Δ*cstA* strains when grown in CDMOBG with 50 mM pyruvate (Figure 6D). This would be consistent with previous data showing that *CD630_23730* is repressed by CcpA in TY medium, but not in TY medium supplemented with glucose (Antunes et al, 2012). The expression of additional pyruvate importers in the presence of preferred carbon sources, such as glucose are in agreement with our findings and those in other bacteria, such as *E. coli* (Ogasawara et al., 2019).

In the absence of glucose, our data indicate that pyruvate uptake leading to biofilm formation is dependent on CstA. However, in the presence of glucose, biofilm formation is dependent on glucose metabolism followed by a critical metabolic shift requiring pyruvate uptake by CstA and other importers. In support of this, we observed that the Δ*ccpA*Δ*cstA* strain used the same amount of extracellular pyruvate as the parental strain after 24 h, but pyruvate uptake in the Δ*ccpA*Δ*cstA* strain was 5 times less (1-fold decrease in extracellular pyruvate) than the parental strain (5-fold decrease in extracellular pyruvate) between 24-48h (Figure 6E). This suggests the 24h-48h period is when the critical shift in metabolism occurs. In support, the Δ*ccpA* and Δ*cstA* that formed biofilms also had ∼10-fold and ∼2-3-fold decreases, respectively, in extracellular pyruvate levels between 24-48h. Taken together, active pyruvate uptake in stationary phase, a process that is partially CstA-dependent, drives a metabolic shift in *C. difficile* and leads to biofilm formation. These data confirm the importance of the role of pyruvate uptake in biofilm formation. This lifestyle switch can be driven by efficient pyruvate uptake and/or carbon metabolism.

## Discussion

We previously identified DOC as an inducer of biofilm formation by *C. difficile*. DOC-induced biofilm took more than 24 h to form, suggesting the specifics steps and pathways controlling the shift from planktonic to biofilm could be elucidated. In this study, our objectives were to identify and characterize key molecules and events required for DOC-induced biofilm formation. Using a combination of time-course transcriptomics and deletion strains, we identified several metabolic processes as a key drivers of DOC-induced biofilm formation. We also show that DOC-induced biofilm formation does not require quorum sensing and surface proteins previously associated with biofilm formation in the absence of DOC. However, the T4aP machinery encoded by the *pilA1* cluster is required for biofilm formation. Based on these findings and the analysis of culture supernatants, we demonstrated the importance of extracellular pyruvate and its integration for biofilm formation.

In some bacteria, pyruvate is known to be excreted during overflow metabolism (Tomlinson and Hochstein, 1972; Ruby and Nealson, 1977; Charbonnier et al. 2017; van den Esker et al. 2017). In our study, *C. difficile* is grown in BHISG and this medium provides an environment rich in proteins with an excess of glucose, which is suited for overflow metabolism (Sonenshein, 2007). Based on our transcriptomic analysis, DOC has a profound impact on metabolism-associated genes, specifically glycolysis (Supplementary Figure S2). The presence of DOC induces a metabolic stress that probably leads to overflow metabolism between inoculation and 14h, the excretion of pyruvate and, once glucose is exhausted, biofilm formation (see the proposed model in Figure 7). The mechanism by which pyruvate is excreted remains to be elucidated in bacteria and this appears to be independent of pyruvate importers (Gasperotti et al., 2020). Furthermore, the absence of the metabolic regulatory factors SigL, CcpA, and CodY resulted in decreased biofilm formation as shown in this work, and Dubois et al. (2019). Specifically, these regulators are important to control flux between different metabolic pathways and help *C. difficile* transition to different sources of energy (Dineen et al, 2010; Antunes et al., 2012; Soutourina et al, 2020). Interestingly, the *sigL* inactivated strain was previously shown to require glucose for optimal growth and does not excrete pyruvate (Dubois et al., 2016; Soutourina et al, 2020). Given that the *sigL* inactivated strain did not form biofilm, this is consistent with our hypothesis that pyruvate excretion drives DOC-induced biofilm formation. Consumption of a preferred carbon source (e.g. glucose) is also important since a strain lacking *ptsI* was unable to form biofilms in the presence of glucose. In addition to glucose, other sugars used by *C. difficile* that require their own uptake systems, such as Neu5Ac and the NanEAT system, might also potentiate the metabolic shift required for biofilm formation. Taken together, our data indicate that in presence of DOC, *C. difficile* must control its metabolic activity to ensure overflow metabolism is activated to promote stationary phase survival and biofilm formation (Figure 7).

**Figure 7.**
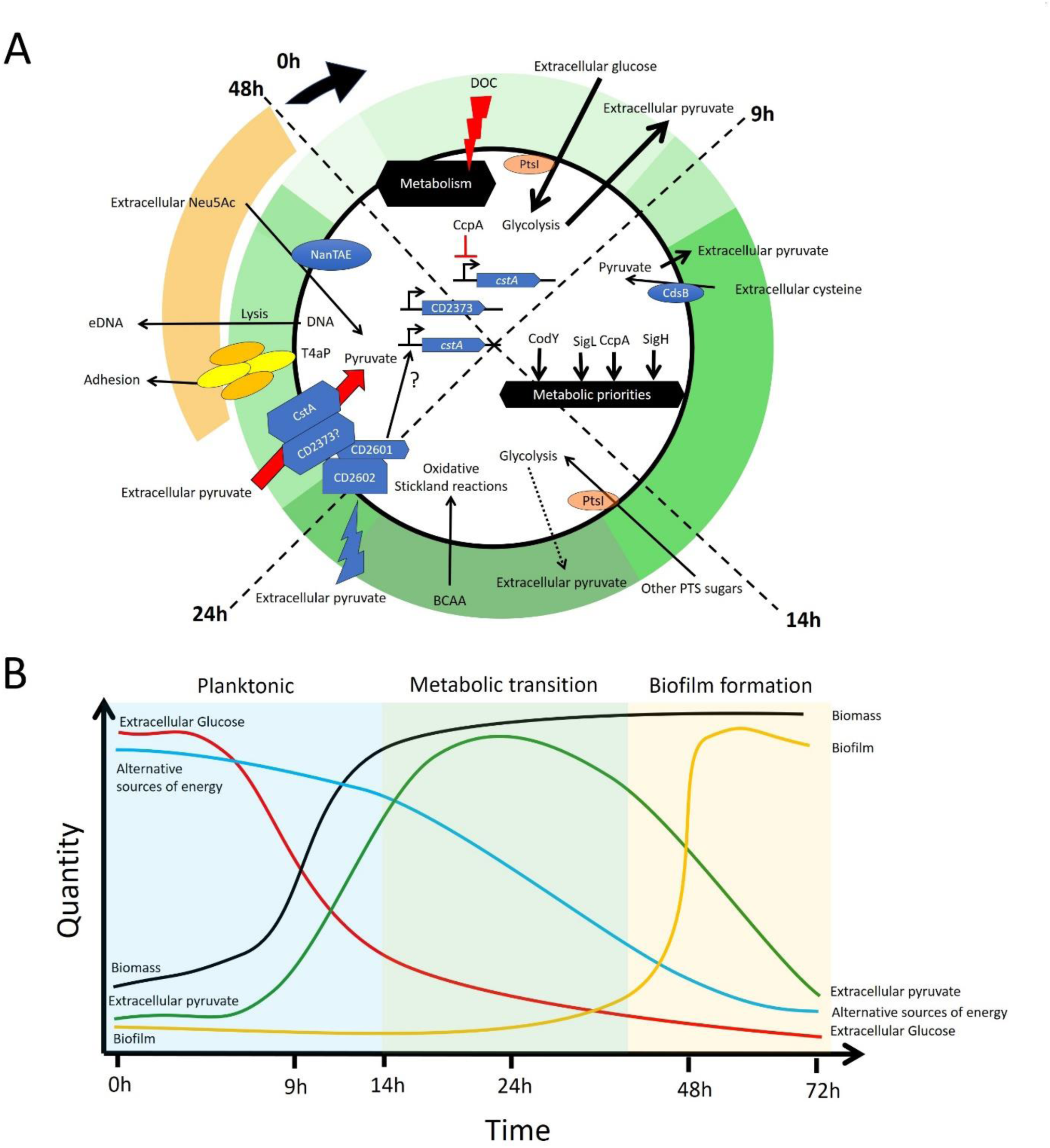
Proposed model of cellular processes leading to DOC-induced biofilm formation. (A) Important cellular processes that contribute to metabolic adaptation and biofilm formation are highlighted for each time interval. Green outer ring indicates the concentration of extracellular pyruvate; darker shades of green are indicative of increased pyruvate concentrations (i.e. light green is the lowest concentration and dark green is the highest concentration). The straw-colored arc indicates when *C. difficile* forms a biofilm. Arrows indicate cellular processes that occur, and the thickness of the arrows indicates how essential they are for biofilm formation (i.e. dashed arrows: least essential, thick, full arrows: absolutely required). (B) Line graph showing hypothetical model of the changes in the quantity of extracellular nutrients, biomass, and biofilm over time. Briefly, during exponential phase (light blue shading), the PTS will import glucose leading to excretion of pyruvate. Glucose is preferentially used until the bacteria enter stationary phase (light green shading) after approximately 14 h of growth. After entry into stationary phase, there is a metabolic shift driven by CcpA, SigH, SigL and CodY, and cells start using alternative sources of energy. As *C. difficile* progresses through stationary phase, it will sense extracellular pyruvate via CD630_26010-26020 TCS and will start actively importing extracellular pyruvate via CstA and other importers such as CD630_23730. The use of extracellular pyruvate maintains *C. difficile* viability during the stationary phase and the type IVa pili (T4aP) machinery will start assembly pili to enhance adherence. A sub-population of the cells will undergo lysis contributing eDNA for the biofilm matrix and, as time passes, the biofilm biomass will increase reaching its peak at approximately 48 h (biofilm phase indicated in yellow).

The importance of overflow metabolism and the excretion of pyruvate likely explain why casein hydrolysate supported biofilm formation whereas casamino acids and specific essential amino acids did not. Casein hydrolysate is a richer source of small peptides and amino acids than casamino acids or purified amino acids. This explains the additive effect of BCAA supplementation on biofilm formation as adding BCAAs may replace the BCAA synthesized as end-products of pyruvate metabolism. BCAAs are likely used in the oxidative Stickland reaction to produce energy during late stationary phase to promote biofilm formation. Overall, it appears that our semi-defined medium was optimized to support overflow metabolism and biofilm formation by *C. difficile*.

The key role of extracellular pyruvate for biofilm formation is not limited to *C. difficile*. Recently, it was demonstrated that *Staphylococcus aureus* requires the presence of extracellular pyruvate to form and maintain a biofilm (Goodwine et al., 2019). In *Streptococcus mutans*, pyruvate improves stationary phase survival and protects against microbiota-generated oxidative stresses in a density- and CcpA-dependent fashion (Ahn et al., 2019; Ishkov et al., 2020; Redanz et al., 2020). Furthermore, pyruvate fermentation is important for *Pseudomonas aeruginosa* microcolony formation and long-term survival in anaerobic environments (Eschbach et al., 2004; Petrova et al. 2012). In this case, lactate is oxidized to pyruvate by the cells in the oxygen-rich top layer of the biofilm and the secreted pyruvate is then converted to acetate by the cells in the anoxic lower layers (Eschbach et al., 2004; Petrova et al. 2012). This metabolic cooperation within the biofilm community is crucial, as it provides a means for the cells to produce enough ATP to survive, but not grow, in the nutrient poor environment of the deep layer of the biofilm (Stewart et al. 2019). Unlike *P. aeruginosa*, *B. subtilis* uses a lactate to promote optimal biofilm formation (Chai et al. 2009). However, *C. difficile* is a strict anaerobe and does not need to switch from an aerobic based metabolism to an anaerobic-based metabolism. We must consider that the cells in the deep layers of the biofilm have limited resources. Our transcriptomic analysis supports that biofilm cells reprioritise their metabolism, as we observed major shifts in the expression of several genes associated with metabolic pathways at 48 h. In our transcriptional analysis, genes associated with fermentation were up-regulated (Figure 2) at 24 h but not 48 h. We also found evidence that butyric acid and lactic acid production occurs between 24 h and 48 h (Supplementary Figure S6B). Therefore, *C. difficile* probably starts using extracellular pyruvate after 24 h of incubation, which helps long-term survival of the cells during stationary phase. In support of this, *Haemophilus influenzae* uses pyruvate as a pivotal point in metabolic adaptation for biofilm cells (Harrison et al., 2019). This reprioritization is critical for metabolic adaptation and long-term survival of biofilm cells.

A recent study provides some interesting insight into biofilm formation that is dependent on eDNA (Yu et al. 2018). In this study, sub-inhibitory concentrations of antibiotics that target the cell envelope enhance biofilm formation by increasing the amount of DNA release from lysing cells without affecting the overall viability of the population. From their data, the authors build a mathematical model that accounts for cell lysis and death, viability, growth and aggregation or binding provided by eDNA. Using this model, we can understand the importance of metabolism in the induction of biofilm in the presence of DOC or pyruvate given that the DOC-induced biofilm formed by *C. difficile* is eDNA-dependent (Dubois et al., 2019). In our proposed model, extracellular pyruvate improves long term viability of *C. difficile* during stationary phase which compensates for autolysis (Figure 7). As DNA is released by lysis, viable cells start aggregating, which may involve the T4aP, as time passes, the biomass increases and *C. difficile* adapts its metabolism to this sedentary lifestyle. Overall, our growth conditions allow *C. difficile* to pass the “biofilm threshold” and any disturbance to the metabolism (SigL, CcpA, CodY, PtsI), lysis (Cwp19), binding or aggregation (eDNA, T4aP) would prevent *C. difficile* from crossing this threshold (Figure 7).

Extracellular pyruvate is also produced by the gut microbiota and provides protection against colonization by *Salmonella* (Morita et al. 2019). In this case, pyruvate concentrations were higher in the luminal content of specific pathogen free (SPF) mice than those of germ-free mice or SPF mice treated with oral vancomycin or neomycin. Furthermore, colonization by the gut commensal bacterium *Lactobacillus helveticus*, which excretes high concentrations of pyruvate *in vitro*, increased pyruvate concentrations in SPF mice. Therefore, it is possible that *C. difficile* encounters extracellular pyruvate during colonization of the intestinal tract containing a normal microbiota or when the microbiota is restored after antibiotic therapy.

Depending on the composition of the microbiota, *C. difficile* could encounter favourable conditions that would include sub-inhibitory concentrations of DOC, and availability of specific amino acids, mucus-derived sugars, and pyruvate (Abbas and Zackular, 2020; Girinathan et al., 2020; Pereira et al., 2020). Under these favourable conditions, sub-inhibitory concentration of DOC would trigger a metabolic adaptation in *C. difficile* to use the available metabolites produced by the microbiota. Interestingly, the molecules that trigger biofilm formation also repress sporulation (DOC; Dubois et al 2019) and toxin production (DOC and pyruvate; Dubois et al. 2016; 2019). Overall, conditions that are favourable for long-term colonization and biofilm formation promote *C. difficile* persistence rather than sporulation or virulence. Any disturbance to this balance, such as the disappearance of inhibitory molecules (e.g. DOC), increase nutrient availability and change in SCFA profiles, could trigger blooms of *C. difficile* as observed recently (VanInsberghe et al. 2020). Furthermore, other signals, such as change in amino acids availability or increase SCFA production, could prevent *C. difficile* from entering a biofilm or persistence state and promote sporulation. These hypotheses are supported by recent *in silico* modeling of the metabolism in the context of sporulation and virulence where each context has distinct metabolic intake and efflux (Jenior et al., 2020).

In summary, we have identified key determinants of DOC-induced biofilm by *C. difficile*. These determinants are unique to DOC-induced biofilm formation suggesting a distinct mechanism. These includes regulator of lifestyle and metabolism and T4aP but the most interesting finding is the importance of extracellular pyruvate and its integration in promoting biofilm formation. Early in our growth conditions (before 14h), pyruvate is probably excreted as a result of overflow metabolism and, as *C. difficile* progresses from exponential to stationary phase, extracellular pyruvate is imported using CstA and other pyruvate importers (Figure 7). This prevents rapid cell death and allows *C. difficile* to generate eDNA through autolysis and pass the “biofilm threshold”. Interestingly, extracellular pyruvate is produced in the gut by commensal bacteria and this could act as a source of pyruvate for *C. difficile*. In conclusion, extracellular pyruvate in the presence of other microbial metabolite could act as a key molecule driving *C. difficile* persistence in the intestinal tract or in response to DOC.

## Methods

### Bacterial Strains and culture conditions

Bacterial strains and plasmids used in this study are listed in Supplementary Table 3. *E. coli* strains were grown in LB broth with chloramphenicol (15 μg/ml). *C. difficile* strains were grown anaerobically (5% H_2_, 5% CO_2_, 90% N_2_) in BHISG (BHI supplement with 0.5% (w/v) yeast extract, 0.01mg/mL cysteine and 100 mM glucose). Additionally, 10 ng/ml of anhydrotetracycline (Atc) was used to induce the *P_tet_* promoter of pRPF185 vector derivatives in *C. difficile*.

The final composition of CDMOB is as follow: Oxoid casein hydrolysate (10 mg/mL), L-Tryptophane (0.5 mg/mL), L-Cysteine (0.01 mg/mL), L-Leucine (0.0033 mg/mL), L-Isoleucine 0.0033 mg/mL), L-Valine (0.0033 mg/mL), Na_2_HPO_4_ (5 mg/mL), NaHCO_3_ (5 mg/mL), KH_2_PO_4_ (0.9 mg/mL) NaCl (0.9 mg/mL), (NH_4_)_2_SO4 (0.04 mg/mL), CaCl_2_·2H_2_O 0,026 MgCl_2_·6H_2_O (0,02 mg/mL), MnCl_2_·4H_2_O (0,01 mg/mL), CoCl26H2O (0.001 mg/mL) FeSO_4_·7 H_2_O (0.004 mg/mL) D-biotine (0.001 mg/mL), calcium-D-panthothenate (0.001 mg/mL) and pyridoxine (0.0001 mg/mL). The desired sugars and/or DOC were added, as necessary.

### Biofilm assays

Overnight cultures of *C. difficile* were diluted 1/100 into the desired medium (BHIS or CDMOB) containing the desired supplements (100 mM glucose, 240 DOC and/or 50 mM or 100 mM sodium pyruvate) and 1 ml of the dilution was aliquoted in each well of a 24-well polystyrene tissue culture-treated plates (Costar, USA). Plates were incubated at 37°C in an anaerobic environment for 48h. Biofilm biomass was measured using established methods (Dubois et al 2019). Briefly, spent media was removed by inverting the plate and wells were washed twice by pipetting phosphate-buffered saline (PBS) at 45° angle. Biofilms were air dried and stained with crystal violet (CV; 0.2% w/v) for 2 min. CV was removed by inversion; wells were washed twice with PBS then air-dried. Dye bound to the biofilm biomass was solubilized by adding 1 ml of a 75% ethanol solution and the absorbance, corresponding to the biofilm biomass, was measured at a λ_600nm_ with a plate reader (Promega GloMax Explorer). Sterile medium was used as a negative control and a blank for the assays.

### RNA isolation and quantitative reverse-transcriptase PCR

A 24-well plate was used to produce one replicate for one condition. At 9 h, 14 h and 24 h, the total bacterial population was collected, and cells were harvested by centrifugation (10 min, 4000 × g, 4°C). The pellet was frozen (−80°C) until used. For the 48h biofilm sample, the supernatant was removed by inverting the plate, the biofilm was washed twice and resuspended in 20 mL of PBS. The recovered biofilm cells were centrifuged and the pellet was frozen until RNA was extracted. Total RNA was extracted from cell pellets as previously described (Saujet et al. 2013). cDNA synthesis and qRT-PCR were carried as described before (Saujet et al. 2013) using primers listed in Supplementary Table 4.

### Whole transcriptome sequencing and analysis

Transcriptomic analysis for each condition was performed using 3 independent RNA preparations using methods described before (Dubois et al., 2019). Briefly, the RNA samples were first treated using Epicenter Bacterial Ribo-Zero kit. This depleted rRNA fraction was used to construct cDNA libraries using TruSeq Stranded Total RNA sample prep kit (Illumina). Libraries were then sequenced by Illumina HiSeq2500 sequencer. Cleaned sequences were aligned to the reannotated *C. difficile* strain 630 (Monot et al., 2011) for the mapping of the sequences using Bowtie 2 (Version 2.1.0). DEseq2 (version 1.8.3) was used to perform normalization and differential analysis using the 9h time point values as a reference for reporting the expression data of the 14 h, 24 h and 48 h. Genes were considered differentially expressed if the fold changes were ≥ Log_2_ 1.5 and their adjusted p-value was ≤ 0.05.

### Gene deletion inactivation and complementation in *C. difficile*

Gene deletions were carried as described in Peltier et al (2020). Briefly, regions upstream and downstream of the gene of interest were PCR-amplified using primer pairs listed in Supplementary Table 4. PCR fragments and linearized pDIA6754 (Peltier et al. 2020) were then mixed and assembled using IVA cloning (Garcià-Nafria et al. 2015) or Gibson assembly and transformed by heat shock into *E. coli* NEB 10β. Constructions were verified by sequencing and the selected plasmid were introduced into *E. coli* HB101 (RP4). Plasmids were transferred by conjugation into the desired *C. difficile* strains and deletion mutants were obtained using counter-selection described elsewhere (Peltier et al., 2020).

To complement the *ptsI*-deletion strain, the *ptsI* gene with its RBS was PCR amplified using appropriate primers (Supplementary Table 4) and inserted into the *Sac*I and *BamH*I restriction sites of pRPF185 (Soutourina et al. 2013) using IVA cloning to generate plasmid pDIA6996. To complement the *cdsB* deletion strain, *cdsB* and its promoter were amplified by PCR using the primers listed in Supplementary Table 4 and inserted in the restriction site BamHI and XhoI of pMTL84121 (Heap et al. 2009) using IVA cloning to generate plasmid pDIA6997. Both plasmids were then transferred by conjugation into the desired strains, yielding strains CDIP1169 and CDIP1170 respectively.

### Enzymatic dispersion of biofilms

Biofilm dispersion experiments were performed as described previously (Dubois et al. 2019). Briefly, biofilms were grown in CDMOBG with 240 µM DOC as described above and, after 48 h, 50 µl of a DNase I solution (500 µg/ml in water), 50 µl of a proteinase K solution (500 µg/ml in water) or 50 µl of fresh 800 mM NaIO4 in water (for a final concentration of 40 mM) was added directly to the biofilms. Control wells were treated with 50 µl water. Wells were treated under anaerobic conditions at 37°C for 1h with DNase I and proteinase K or for 2h with NaIO4. Biofilms were then washed, stained, and quantified as described above.

### Biofilm matrix analysis

The biofilm matrix was harvested and purified as described previously (Dubois et al. 2019). Briefly, biofilms were grown as described above, washed twice with PBS and resuspended in 1.5 M NaCl (12 wells/mL). The biofilm suspension was then centrifuged (8 000 × g for 10 min) and the supernatant was collected and stored at -20°C. A fraction of the matrix was then treated with DNase I (25 µg) and Proteinase K (25 µg) for 1 h at 37°C. Samples were then analysed by agarose gel electrophoresis or SDS-PAGE. DNA was stained with ethidium bromide, proteins with Coomassie blue, and glycol-proteins and the DNase/Proteinase K treated matrix samples with the Pro-Q Emerald 300 glycoprotein stain (ThermoFisher).

### Gas Phase Chromatography

*C. difficile* 630Δ*erm* was grown in BHISG or BHISG supplemented with 240 µM DOC in 24 well plates. After 24 h or 48 h, 12 mL of culture was recovered and cells were removed by centrifugation (10 min, 4000 × g, 4°C). The supernatants were recovered and stored at -20°C for future use. Volatile and non-volatile fatty acids composition was determined and quantified using a Gas Chromatograph (Model CP3380, Varian Inc., United States) as previously described (Carlier and Sellier, 1989). For control purposes, sterile medium was used to determine the initial composition of the fatty acids in the medium.

### Treatment with pyruvate dehydrogenase

The pyruvate depletion assay was adapted from Goodwine et al. (2019). Briefly, biofilms were prepared in CDMOBG with 240 µM DOC as described above. After 24 h, 250 µL containing 20 mU of pyruvate dehydrogenase and cofactors (2mM CoA, 2 mM ß-NAD^+^, 20 µM thiamine pyrophosphate and 50 µM MgSO_4_) was added to the biofilm. For control purposes, pyruvate dehydrogenase was heat inactivated at 100°C for 10 min and mixed with its cofactors. A cofactor-only control and a non-treated control were also included. The values are reported as a percent of the biofilm formed using the following formula: (Treated biofilm (Enzyme, heat-inactivated enzyme or buffer only)/untreated biofilm) × 100.

### Quantification of pyruvate in culture supernatant

*C. difficile* strains were grown in 1mL of CDMOB with 100 mM pyruvate or CDMOBG with 50 mM pyruvate aliquoted in individual wells of a 24-well plate. After 24 h and 48 h, 1 mL of each sample was recovered and the supernatant was recovered by centrifugation (1 min, 14000 × g). The clarified supernatant was transferred to a new tube and stored -20°C until used. Sterile medium was used as a control to quantify the amount of pyruvate at time 0 h. Pyruvate was quantified using the EnzyChrom^TM^ Pyruvate Assay kit (BioAssay Systems). The values are reported as a percent of the pyruvate remaining in the supernatant calculated with the following formula: (Concentration of pyruvate in culture supernatant/Concentration of pyruvate in sterile medium) × 100.

### Statistical analysis

Biofilm assays, effect of treatment and effect of genetic inactivation or deletion were analysed using a Kruskal-Wallis test followed by an uncorrected Dunn’s test. Effect of pyruvate supplementation was compared and analysed using a two-way ANOVA followed by a Fisher LSD test.

### Data Availability

RNA-Seq data generated in this study are available in the NCBI-GEO with accession no GSE165116. Other data that support the findings of this study are available from the corresponding author upon reasonable request.

## Acknowledgements

We would like to thank Johann Peltier, Imane El Meouche and Laurent Bouillaut for generously providing the CDIP634, *fliC*::*erm*, *rex*::*erm* and *prdB*::*erm* strains. This work was funded by the Institut Pasteur and the “Integrative Biology of Emerging Infectious Diseases” (LabEX IBEID) funded in the framework of the French Government’s “Programme Investissements d’Avenir”. YDNT postdoctoral fellowship was funded by the LabEX IBEID.

## Author Contribution

YDNT, IMV and BD participated in the study design; YDNT, BARD, AH and MO performed experiments; MM provided assistance with the transcriptomic experimental design and analysis; YDNT and BD drafted and edited the manuscript; all authors read and approved the final manuscript.

---

**Figure S1:**
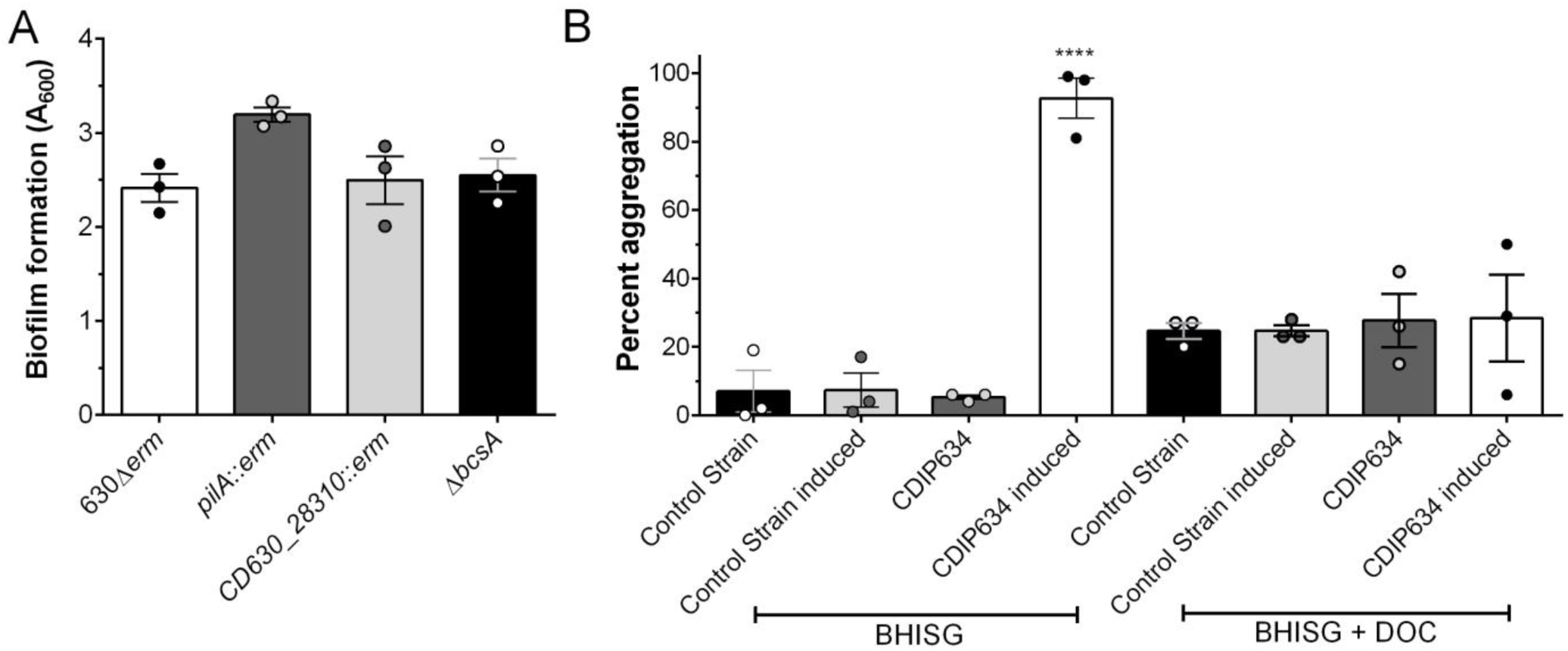
Biofilm formation or aggregation in BHISG supplemented with 240 µM DOC does not require PilA1, CD630_28310, BscA or overproduction of c-di-GMP. (A) Biofilm formation by the *pilA1*::*erm* and the *CD630_630_28310*::erm strains in BHISG with 240 µM DOC. (B) A strain overexpressing CD630_1420 (c-di-GMP overproduction) and a control strain were grown in BHISG or BHISG supplemented with 240 µM DOC in the presence or absence of an inducer in a test tube. Aggregation was assessed after 9h (BHISG) or 24h (BHISG with 240 µM DOC). Percent aggregation = 100 –[100 × (OD_600_ top 1 cm of undisturbed culture / OD_600_ culture after vortexing)]. (C) Biofilm formation by the Δ*bscA* strain in BHISG with 240 µM DOC. Asterisks indicate statistical significance determined with a Kruskal-Wallis test followed by an uncorrected Dunn’s test (**** *p*≤0.001 vs CDIP634). Data shown indicate biological replicates performed on different days, the bars represent the mean and the error bars represent the SEM.

**Figure S2:**
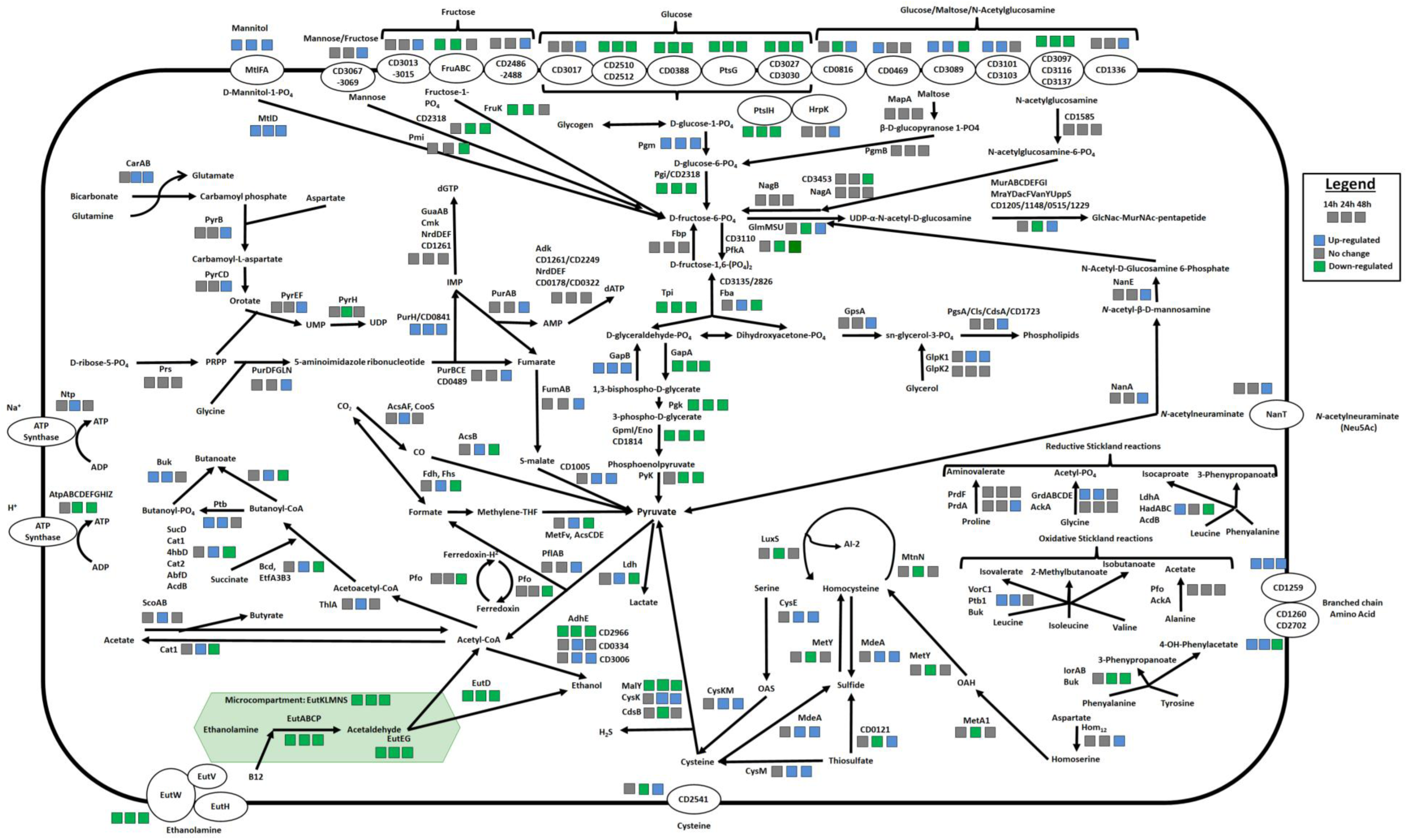
Overview of transcriptomic changes over time in *C. difficile* metabolic pathways during grown in BHISG supplemented with 240 µM of DOC. Cartoon of a bacterial cells with the outside line representing the cell envelope. Cellular processes are place in their predicted cellular location. For example, membrane proteins and transporter are placed on the outside line. Each group of three squares indicate a specific time point (left to right: 14 h, 24 h and 48 h, respectively) and the changes in expression are colour coded (in grey, no changes; blue, up-regulated; or green, down-regulated). The 9 h time point was used as the reference point. Gene ID are used instead of the locus tag to prevent clutter, please see Supplementary data for equivalencies.

**Figure S3:**
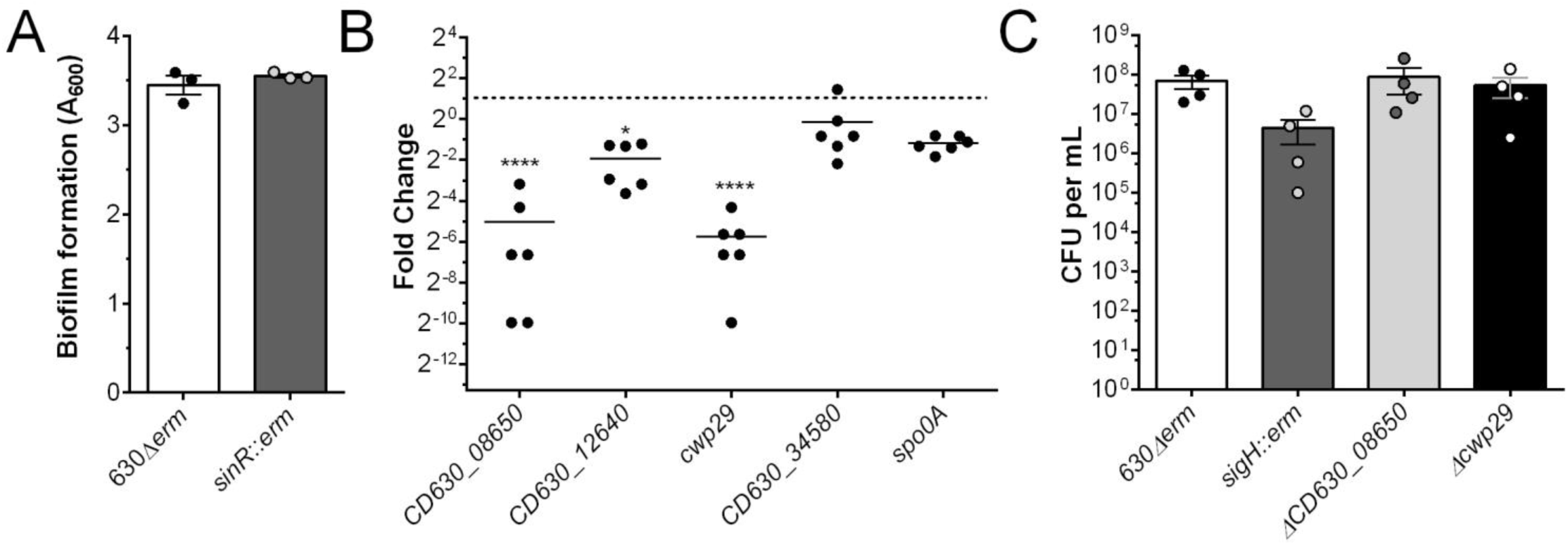
SinR is not required for biofilm formation and CD630_08650 and *cwp29* expression is induced by SigH. (A) Biofilm formation by the *sinR*::*erm* strain in BHISG supplemented with 240 µM DOC. (B) Change in expression of *CD630_08650*, *CD630_12640*, *cwp29* and *CD630_34580* in the *sigH::erm* strain compared to the parental strain after 24 h growth in BHISG supplemented with 240 uM DOC. Expression was normalized to *codY* and *rex*; *spo0A* expression is used as a positive control. The dotted line represents the no changes in expression threshold. (C) Number of viable vegetative cells recovered for the parental, *sigH*::*erm*, Δ*CD630_08650* and Δ*cwp29* strains after 24 h of growth in BHISG supplemented with 240 µM DOC. Asterisks indicate statistical significance determined with a Kruskal-Wallis test followed by an uncorrected Dunn’s test (* *p*≤0.05, **** *p*≤0.0001 vs 630Δ*erm*). Data shown indicate biological replicates performed on different days, the bars represent the mean and the error bars represent the SEM.

**Figure S4.**
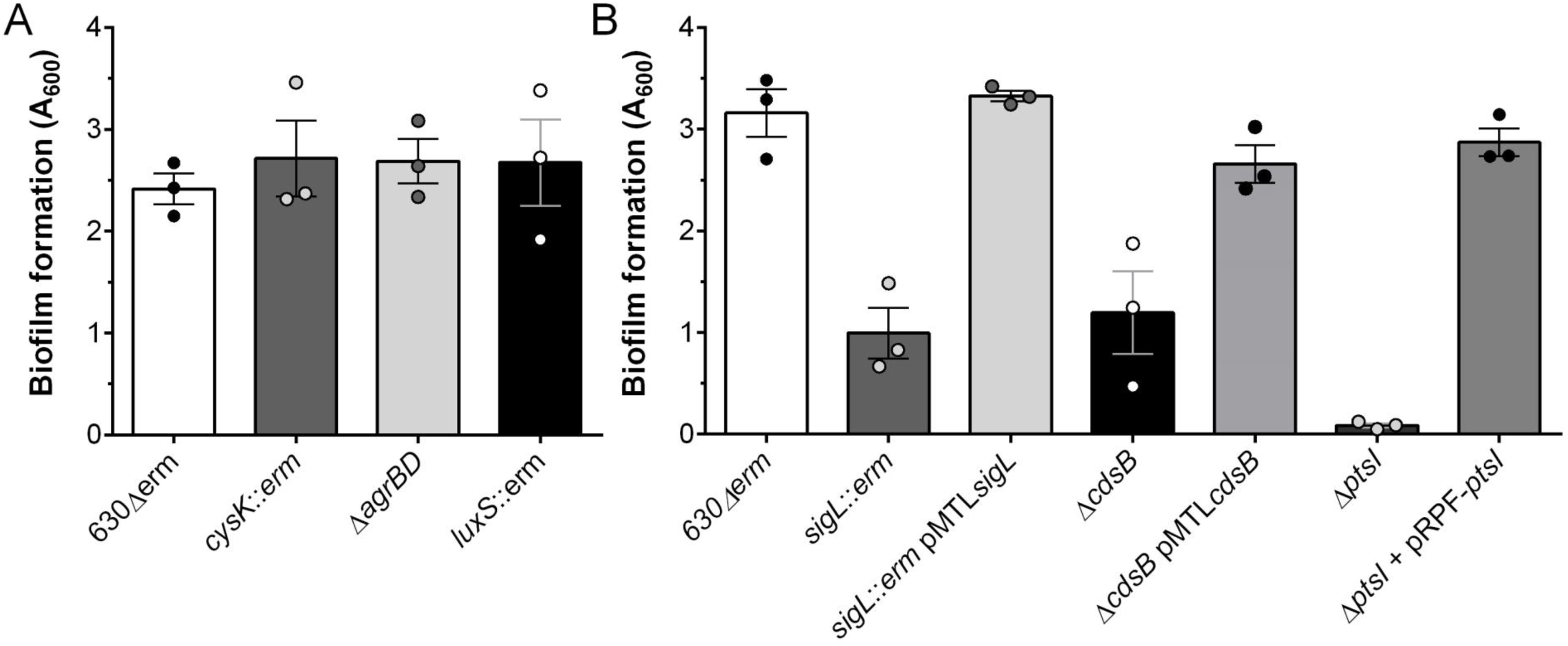
Absence of *cysK*, *luxS* or *agrBD* does not affect biofilm formation and complementation of the *ptsI* or *cdsB* strains restore biofilm formation in BHISG supplemented with 240 µM DOC. Data shown indicate biological replicates performed on different days, the bars represent the mean and the error bars represent the SEM.

**Figure S5:**
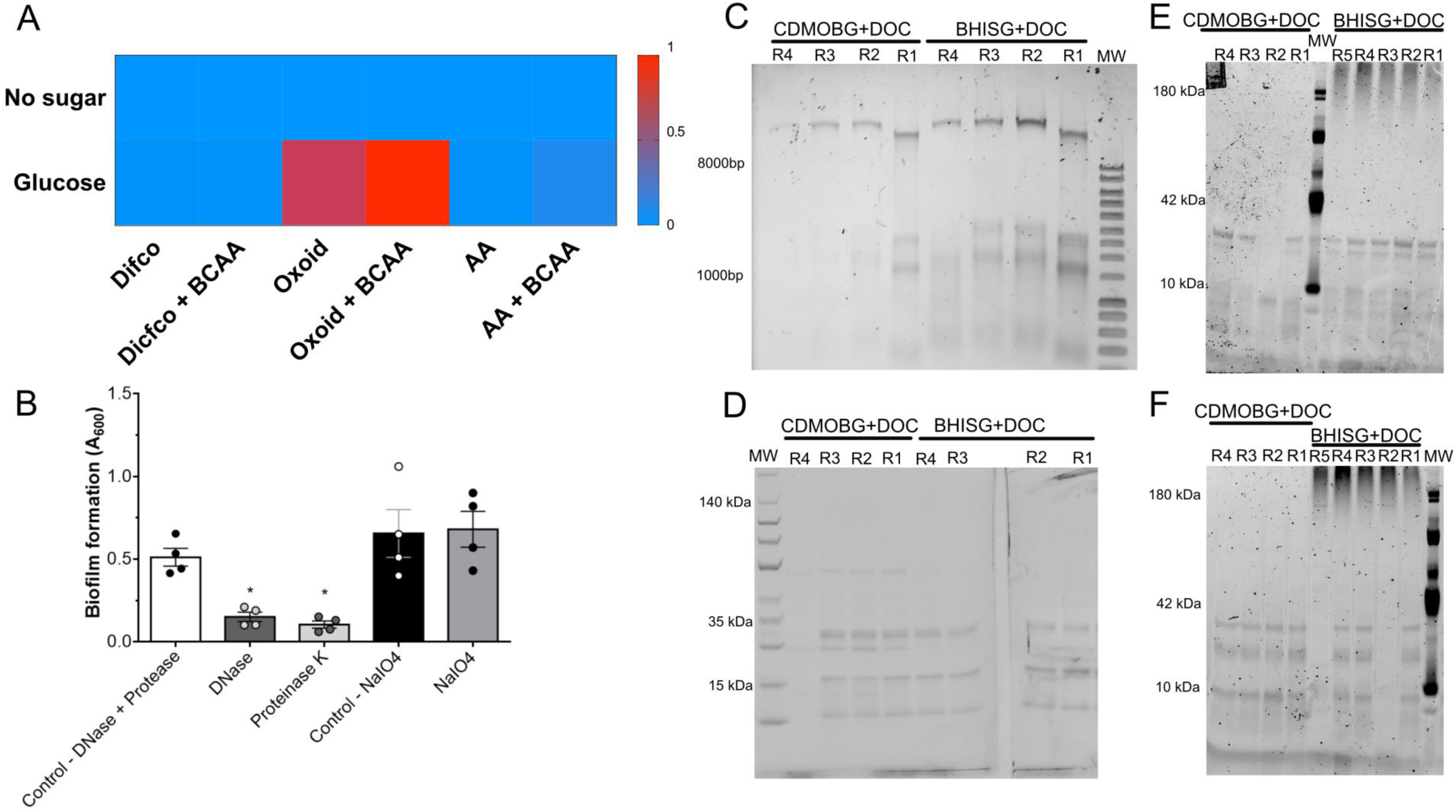
Analysis and optimization of biofilm formation in a semi-defined medium. Heat map for the CDMM optimization with different amino acid sources (A). Enzymatic and chemical dispersion of the preformed biofilm in CDMOBG supplemented with 240 μM DOC (B). Asterisks indicate statistical significance determined with a Kruskal-Wallis test followed by an uncorrected Dunn’s test (* *p*≤0.05 vs WT). Data shown indicate biological replicates performed on different days and the bar represent the mean. Agarose gel electrophoresis confirming the presence of eDNA in the biofilm matrix (C). SDS-PAGE analysis of the biofilm matrix for the presence of proteins (D), glycoproteins (E) and DNase/protease-resistant material (F). DNA was stained with ethidium bromide, proteins with Coomassie blue, and glycol-proteins and DNase/Proteinase K resistant material with the Pro-Q Emerald 300 glycoprotein stain. R1, R2, R3, R4 and R5 indicate the replicate number.

**Figure S6.**
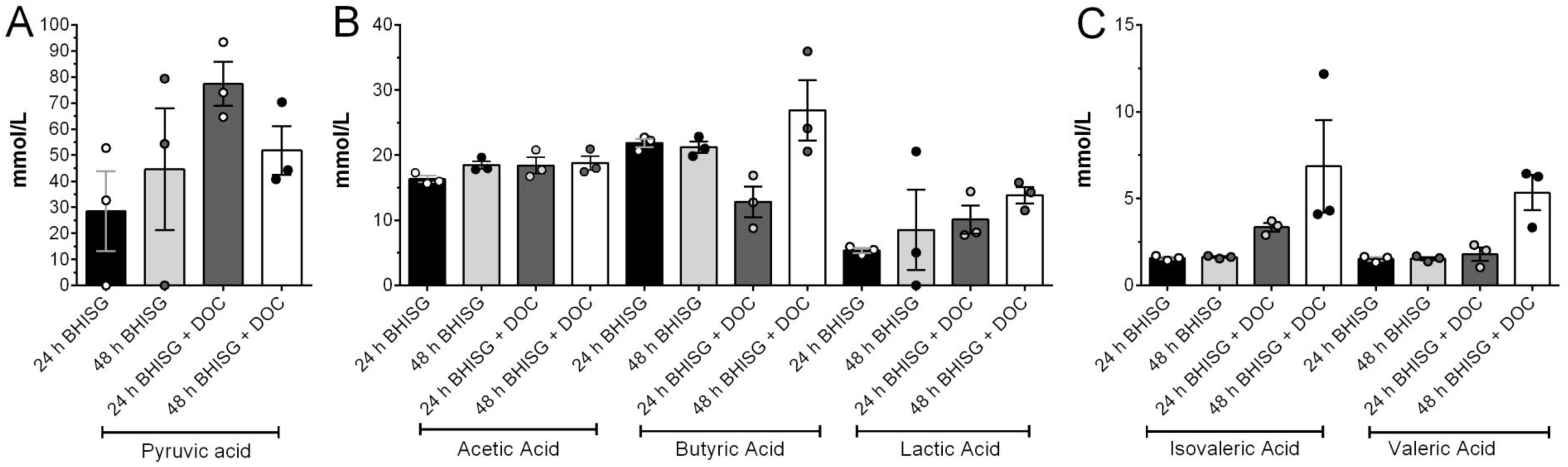
Gas chromate of culture supernatant of *C. difficile* grown in BHISG and BHISG with 240 µM DOC for 24 h or 48 h. To improve visualisation, data was split in based on the amount of each fatty acid detected. (A) pyruvic acid, (B) Acetic, butyric and lactic acid, (C) isovaleric and valeric acid. Data shown indicate biological replicates from supernatant collected on different days, the bar represent the mean and the error bar represent the SEM.

**Table S1:**
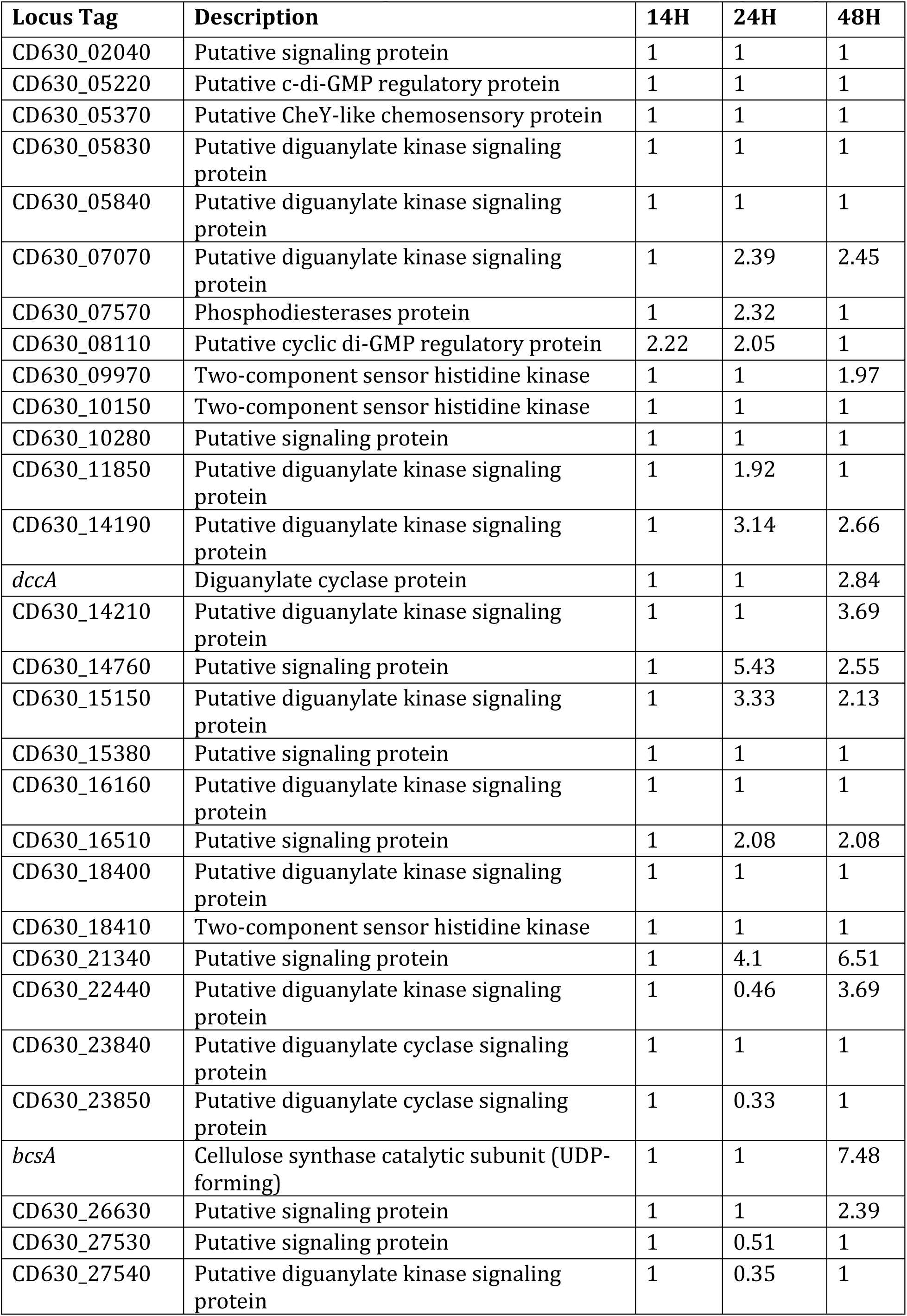
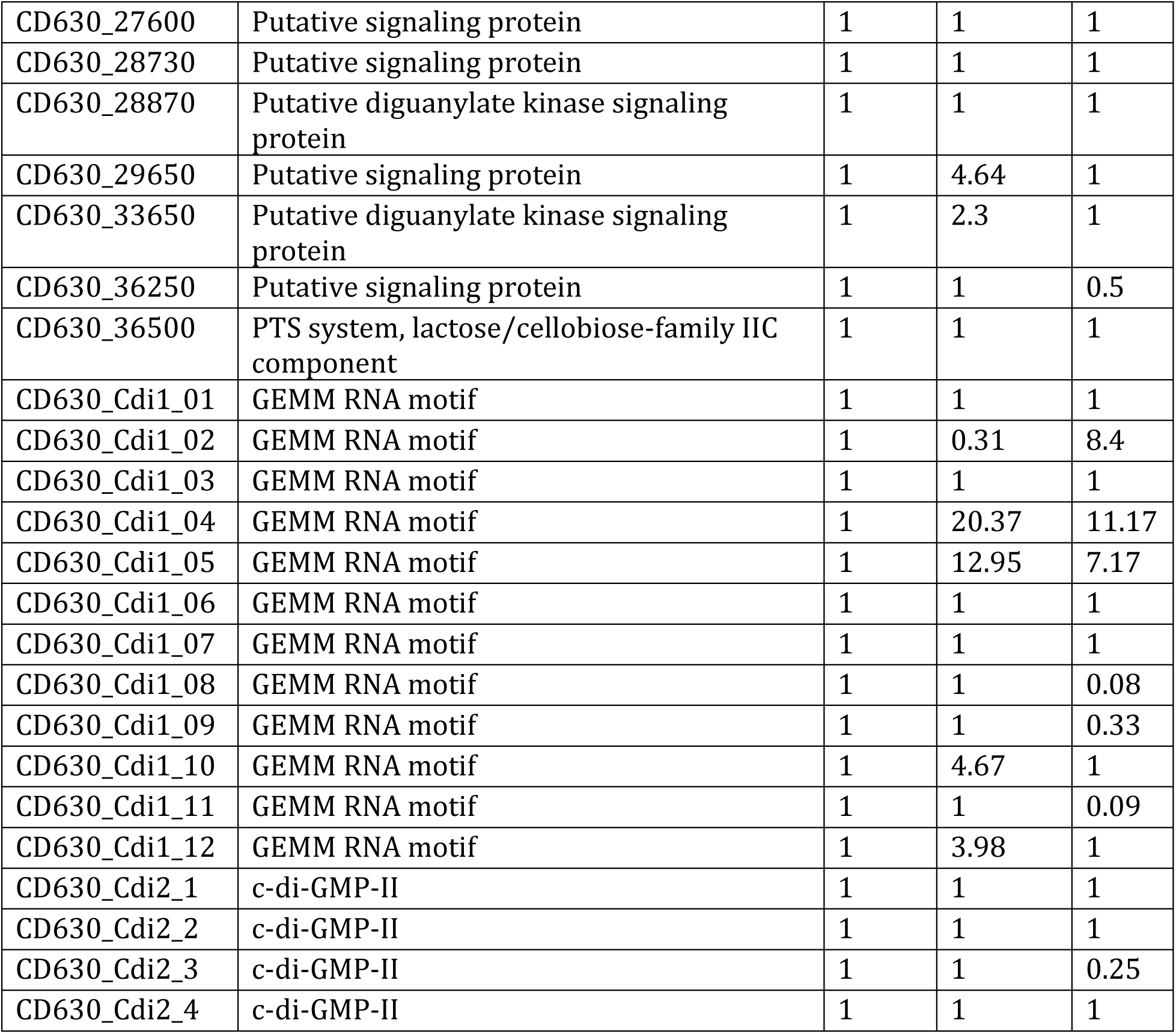
Differential expression of genes associated with c-di-GMP signalling

| **Locus Tag** | **Description** | **14H** | **24H** | **48H** |
| --- | --- | --- | --- | --- |
| CD630_02040 | Putative signaling protein | 1 | 1 | 1 |
| CD630_05220 | Putative c-di-GMP regulatory protein | 1 | 1 | 1 |
| CD630_05370 | Putative CheY-like chemosensory protein | 1 | 1 | 1 |
| CD630_05830 | Putative diguanylate kinase signaling protein | 1 | 1 | 1 |
| CD630_05840 | Putative diguanylate kinase signaling protein | 1 | 1 | 1 |
| CD630_07070 | Putative diguanylate kinase signaling protein | 1 | 2.39 | 2.45 |
| CD630_07570 | Phosphodiesterases protein | 1 | 2.32 | 1 |
| CD630_08110 | Putative cyclic di-GMP regulatory protein | 2.22 | 2.05 | 1 |
| CD630_09970 | Two-component sensor histidine kinase | 1 | 1 | 1.97 |
| CD630_10150 | Two-component sensor histidine kinase | 1 | 1 | 1 |
| CD630_10280 | Putative signaling protein | 1 | 1 | 1 |
| CD630_11850 | Putative diguanylate kinase signaling protein | 1 | 1.92 | 1 |
| CD630_14190 | Putative diguanylate kinase signaling protein | 1 | 3.14 | 2.66 |
| *dccA* | Diguanylate cyclase protein | 1 | 1 | 2.84 |
| CD630_14210 | Putative diguanylate kinase signaling protein | 1 | 1 | 3.69 |
| CD630_14760 | Putative signaling protein | 1 | 5.43 | 2.55 |
| CD630_15150 | Putative diguanylate kinase signaling protein | 1 | 3.33 | 2.13 |
| CD630_15380 | Putative signaling protein | 1 | 1 | 1 |
| CD630_16160 | Putative diguanylate kinase signaling protein | 1 | 1 | 1 |
| CD630_16510 | Putative signaling protein | 1 | 2.08 | 2.08 |
| CD630_18400 | Putative diguanylate kinase signaling protein | 1 | 1 | 1 |
| CD630_18410 | Two-component sensor histidine kinase | 1 | 1 | 1 |
| CD630_21340 | Putative signaling protein | 1 | 4.1 | 6.51 |
| CD630_22440 | Putative diguanylate kinase signaling protein | 1 | 0.46 | 3.69 |
| CD630_23840 | Putative diguanylate cyclase signaling protein | 1 | 1 | 1 |
| CD630_23850 | Putative diguanylate cyclase signaling protein | 1 | 0.33 | 1 |
| *bcsA* | Cellulose synthase catalytic subunit (UDP-forming) | 1 | 1 | 7.48 |
| CD630_26630 | Putative signaling protein | 1 | 1 | 2.39 |
| CD630_27530 | Putative signaling protein | 1 | 0.51 | 1 |
| CD630_27540 | Putative diguanylate kinase signaling protein | 1 | 0.35 | 1 |
| CD630_27600 | Putative signaling protein | 1 | 1 | 1 |
| CD630_28730 | Putative signaling protein | 1 | 1 | 1 |
| CD630_28870 | Putative diguanylate kinase signaling protein | 1 | 1 | 1 |
| CD630_29650 | Putative signaling protein | 1 | 4.64 | 1 |
| CD630_33650 | Putative diguanylate kinase signaling protein | 1 | 2.3 | 1 |
| CD630_36250 | Putative signaling protein | 1 | 1 | 0.5 |
| CD630_36500 | PTS system, lactose/cellobiose-family IIC component | 1 | 1 | 1 |
| CD630_Cdi1_01 | GEMM RNA motif | 1 | 1 | 1 |
| CD630_Cdi1_02 | GEMM RNA motif | 1 | 0.31 | 8.4 |
| CD630_Cdi1_03 | GEMM RNA motif | 1 | 1 | 1 |
| CD630_Cdi1_04 | GEMM RNA motif | 1 | 20.37 | 11.17 |
| CD630_Cdi1_05 | GEMM RNA motif | 1 | 12.95 | 7.17 |
| CD630_Cdi1_06 | GEMM RNA motif | 1 | 1 | 1 |
| CD630_Cdi1_07 | GEMM RNA motif | 1 | 1 | 1 |
| CD630_Cdi1_08 | GEMM RNA motif | 1 | 1 | 0.08 |
| CD630_Cdi1_09 | GEMM RNA motif | 1 | 1 | 0.33 |
| CD630_Cdi1_10 | GEMM RNA motif | 1 | 4.67 | 1 |
| CD630_Cdi1_11 | GEMM RNA motif | 1 | 1 | 0.09 |
| CD630_Cdi1_12 | GEMM RNA motif | 1 | 3.98 | 1 |
| CD630_Cdi2_1 | c-di-GMP-II | 1 | 1 | 1 |
| CD630_Cdi2_2 | c-di-GMP-II | 1 | 1 | 1 |
| CD630_Cdi2_3 | c-di-GMP-II | 1 | 1 | 0.25 |
| CD630_Cdi2_4 | c-di-GMP-II | 1 | 1 | 1 |

**Table S2:** Differential expression of genes with SigH binding box in their promoter

| Gene name | Description | BHISG + 240 μM | | |
| --- | --- | --- | --- | --- |
|  |  | 14h | 24h | 48h |
| *spo0A* | Stage 0 sporulation protein A | 1 | 1 | 1 |
| *CD630_24920* | Histidine kinase associated to Spo0A | 1 | 1 | 1 |
| *Soj* | Sporulation initiation inhibitor | 1 | 1 | 1 |
| *spoIIAA* | Anti-sigma F factor antagonist | 1 | 1 | 0.48 |
| *spoVS* | Stage V sporulation protein S | 1 | 1 | 1 |
| *spoVG* | Regulator required for spore cortex synthesis | 1 | 7.03 | 1 |
| *CD630_36730* | Putative stage 0 sporulation protein, Spo0J-like | 1 | 1 | 1 |
| *CD630_35690* | Sporulation-specific protease | 1 | 1 | 1 |
| *ftsZ* | Cell division protein FtsZ | 1 | 1 | 1 |
| *minC* | Cell division regulator | 1 | 1 | 1.92 |
| *CD630_26240* | Conserved hypothetical protein | 1 | 1 | 1 |
| *sigA2* | Transcription | 1 | 7.24 | 1 |
| *CD630_01420* | Putative RNA-binding protein | 1 | 1 | 1 |
| *CD630_08380* | DNA-binding protein | 2.03 | 1.95 | 1 |
| *gapN* | Glyceraldehyde-3-P dehydrogenase | 1 | 0.41 | 4.79 |
| *glgC* | Glucose-1-P adenylyltransferase | 1 | 11.03 | 3 |
| *fdhF* | Formate dehydrogenase | 3.38 | 4.16 | 0.41 |
| *cobT* | Nicotinate-nucleotide-dimethylbenzimidazole | 2.65 | 1 | 0.21 |
| *CD630_34560* | 5-Formyltetrahydrofolate cycloligase | 1 | 3.2 | 1 |
| *CD630_27380* | Putative cytosine permease | 1 | 1 | 5.45 |
| *CD630_07600* | Putative Ca<sup>2+</sup>/Na<sup>+</sup> antiporter | 1 | 1 | 1 |
| *CD630_27890* | Putative adhesin Cwp66 | 1 | 1 | 1 |
| *CD630_04400* | Putative cell wall-binding protein | 1 | 1 | 1 |
| *cwp29* | Cell surface protein | 1 | 8.3 | 1 |
| *lplA* | Lipoate protein-ligase | 1 | 1 | 1 |
| *CD630_34580* | Membrane protein | 1 | 13.84 | 1 |
| *CD630_28000* | Membrane protein | 1 | 1 | 1 |
| *CD630_22950* | Membrane protein | 1 | 1 | 1 |
| *CD630_15900* | Membrane protein | 1 | 1 | 1 |
| *CD630_08650* | Putative ADP-ribose binding protein | 4.58 | 6.35 | 1 |
| *CD630_24470* | Putative histidine triad (HIT) protein | 1 | 2.36 | 0.49 |
| *CD630_36540* | Putative DNA replication protein | 1 | 1 | 1 |
| *CD630_00220* | Elongation factor G (EF-G) | 6.65 | 82.16 | 1 |
| *CD630_21370* | Ribosome-recycling factor | 1 | 1 | 1 |
| *CD630_19410* | Unknown function | 1 | 1 | 1 |
| *CD630_15431* | Unknown function | 1 | 11.51 | 1 |
| *CD630_12640* | Unknown function | 1 | 9.18 | 1 |
| *CD630_10610* | Unknown function | 1 | 2.29 | 1 |
| *CD630_13170* | Unknown function | 1 | 1 | 3.57 |
| *CD630_16220* | Unknown function | 2.77 | 3.53 | 1 |

**Table S3:** List of strains and plasmids used for qPCR and gene construct

| Strains and plasmids | PCR-ribotype/Genotype | Resistance<sup>1</sup> | Reference |
| --- | --- | --- | --- |
| 630Δerm | 012 | | Hussain et al 2005 |
| CDIP1168 | 630Δerm *sigH::erm* | Erm | Saujet et al. 2011 |
| CDIP3 | 630 Δerm *spo0A::erm* | Erm | Pereira et al. 2013 |
| CDIP1496 | 630Δerm ΔCD630_08650 | | This study |
| CDIP1497 | 630Δerm Δcwp29 (CD630_23050) | | This study |
| CDIP9 | 630Δerm *sinR::erm* (CD630_22140-22150) | Erm | Poquet et al, 2018 |
| *pilA*<sub>1</sub> | 630Δerm *pilA1::erm* (CD630_35130) | Erm | Poquet et al, 2018 |
| CD2831 | 630Δerm *CD630_28310::erm* | Erm | Poquet et al, 2018 |
| CDIP1382 | 630Δerm Δ*pilW* (CD630_23050) | | This study |
| CDIP1443 | 630Δerm ΔT4aP Cluster 1 | | This study |
| CDIP1444 | 630Δerm ΔT4aP Cluster 2 | | This study |
| CDIP1463 | 630Δerm ΔbcsA | | This study |
| CDIP1383 | 630Δerm ΔagrDB (CD630_27491-27500) | | This study |
| CDIP1160 | 630Δerm *fliC::erm* (CD630_02390) | Erm | Iman El Meouche |
| CDIP550 | 630Δerm *sigD::erm* (CD630_02660) | Erm | El Meouche, et al. 2013 |
| CDIP634 | 630Δerm with chromosomal P<sub>tet</sub>-CD630_14200 | | Johann Peltier |
| CDIP1384 | 630Δerm ΔptsI (CD630_27550) | | This study |
| CDIP1697 | CDIP1384 with pDIA6996 | Tm | This study |
| CDIP1385 | 630Δerm ΔhprK (CD630_34090) | | This study |
| CDIP217 | 630Δerm *sigL::erm* (CD630_31760) | Erm | Dubois et al. 2016 |
| CDIP342 | CDIP217 with pDIA6309 | Erm, Tm | Dubois et al. 2016 |
| CDIP1318 | 630Δerm ΔcdsB (CD630_332320) | | This study |
| CDIP1698 | CDIP1318 with pDIA6997 | Tm | This study |
| CDIP540 | 630Δerm *cysK::erm* (CD630_15970) | Erm | Dubois et al. 2016 |
| CDIP1725 | 630Δerm *luxS::erm* (CD630_35980) | Erm | Thomas Dubois |
| CDIP001 | 630Δerm *fur::erm* (CD630_12870) | Erm | Dubois et al. 2016 |
| CDIP1445 | 630Δerm ΔfumAB-CD630_10050 | | This study |
| CDIP1447 | 630Δerm ΔnanAE-CD630_22390 | | This study |
| CDIP537 | 630Δerm *rex::erm* | Erm | Laurent Bouillaut |
| CDIP535 | 630Δerm *prdB::erm* | Erm | Laurent Bouillaut |
| CDIP657 | 630Δerm *CD630_26020::erm* | Erm | Dubois et al. 2016 |
| CDIP1616 | 630Δerm ΔcstA (CD630_26000) | | This study |
| CDIP1335 | 630Δerm ΔccpA | | This study |
| CDIP1629 | 630Δerm ΔccpA ΔcstA | | This study |
| **Plasmids** |  |  |  |
| pDIA6103 | pRPF185ΔgusA | Cm/Tm | Soutourina et al, 2013 |
| pDIA5929 | pMTL84121 | Cm/Tm | Heap et al., 2009 |
| pDIA6753 | pMSR | Cm/Tm | Peltier et al. 2020 |
| pDIA6794 | pMSR-Δ*agrBD* | Cm/Tm | This study |
| pDIA6795 | pMSR-Δ*pilW* | Cm/Tm | This study |
| pDIA6796 | pMSR-Δ*cdsB* | Cm/Tm | This study |
| pDIA6797 | pMSR-Δ*hprK* | Cm/Tm | This study |
| pDIA6821 | pMSR-Δ*ptsI* | Cm/Tm | This study |
| pDIA6939 | pMSR-Δ T4aP Cluster 1 | Cm/Tm | This study |
| pDIA6940 | pMSR-Δ T4aP Cluster 2 | Cm/Tm | This study |
| pDIA6941 | pMSR-Δ*fumAB-CD1005* | Cm/Tm | This study |
| pDIA6945 | pMSR-Δ*bcsA* | Cm/Tm | This study |
| pDIA7043 | pMSR-Δ*cstA* | Cm/Tm | This study |
| pDIA6820 | pMSR-Δ*ccpA* | Cm/Tm | This study |
| pDIA6979 | pMSR-Δ*CD0865* | Cm/Tm | This study |
| pDIA6980 | pMSR-Δ*cwp29* | Cm/Tm | This study |
| pDIA6996 | pRPF185-*ptsI* | Cm/Tm | This study |
| pDIA6997 | pMTL84121-*cdsB* | Cm/Tm | This study |

*Cm* : chloramphenicol; *Tm* : Thiamphenicol; *Erm* : Erythromycin

**Table S4:**
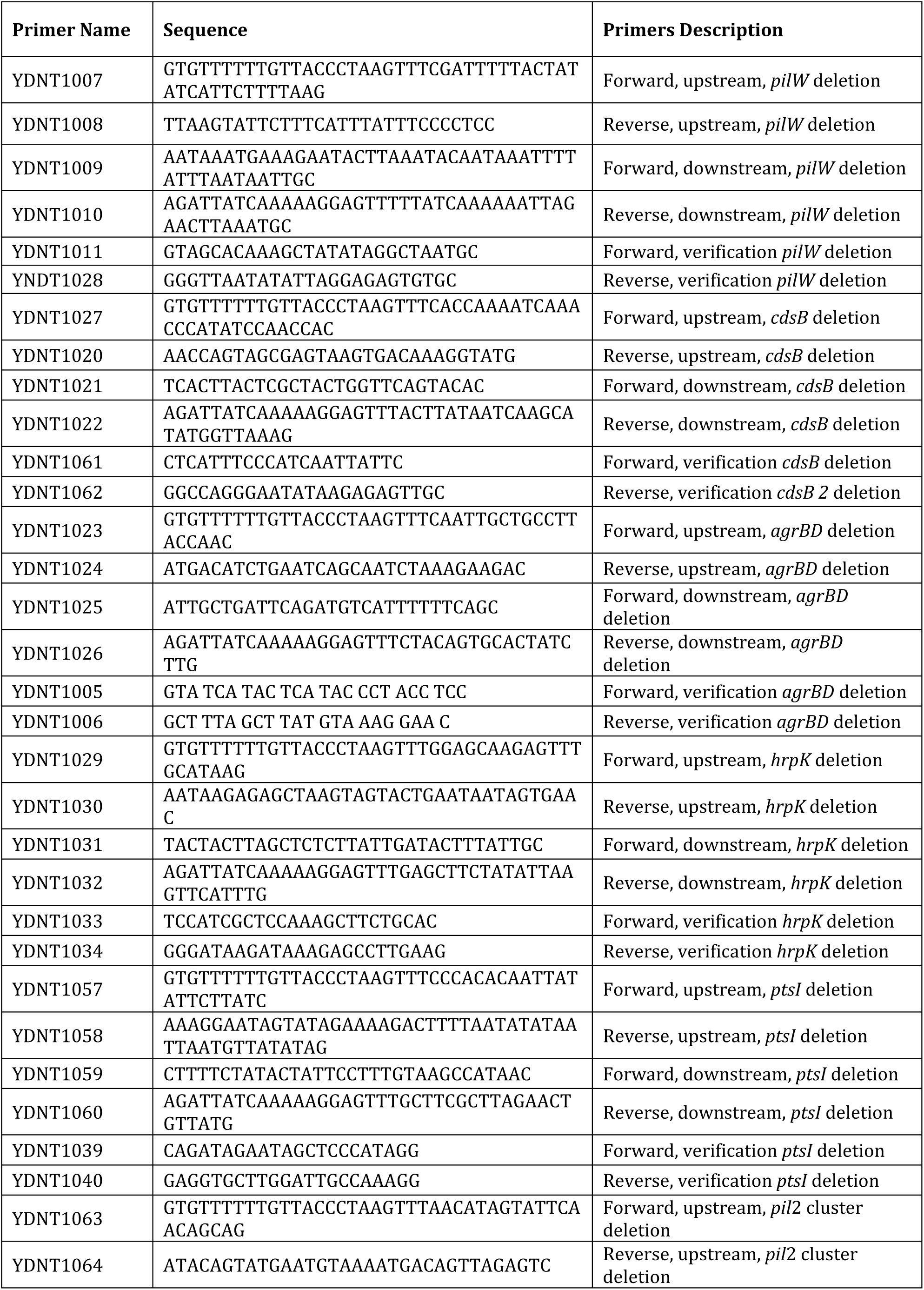
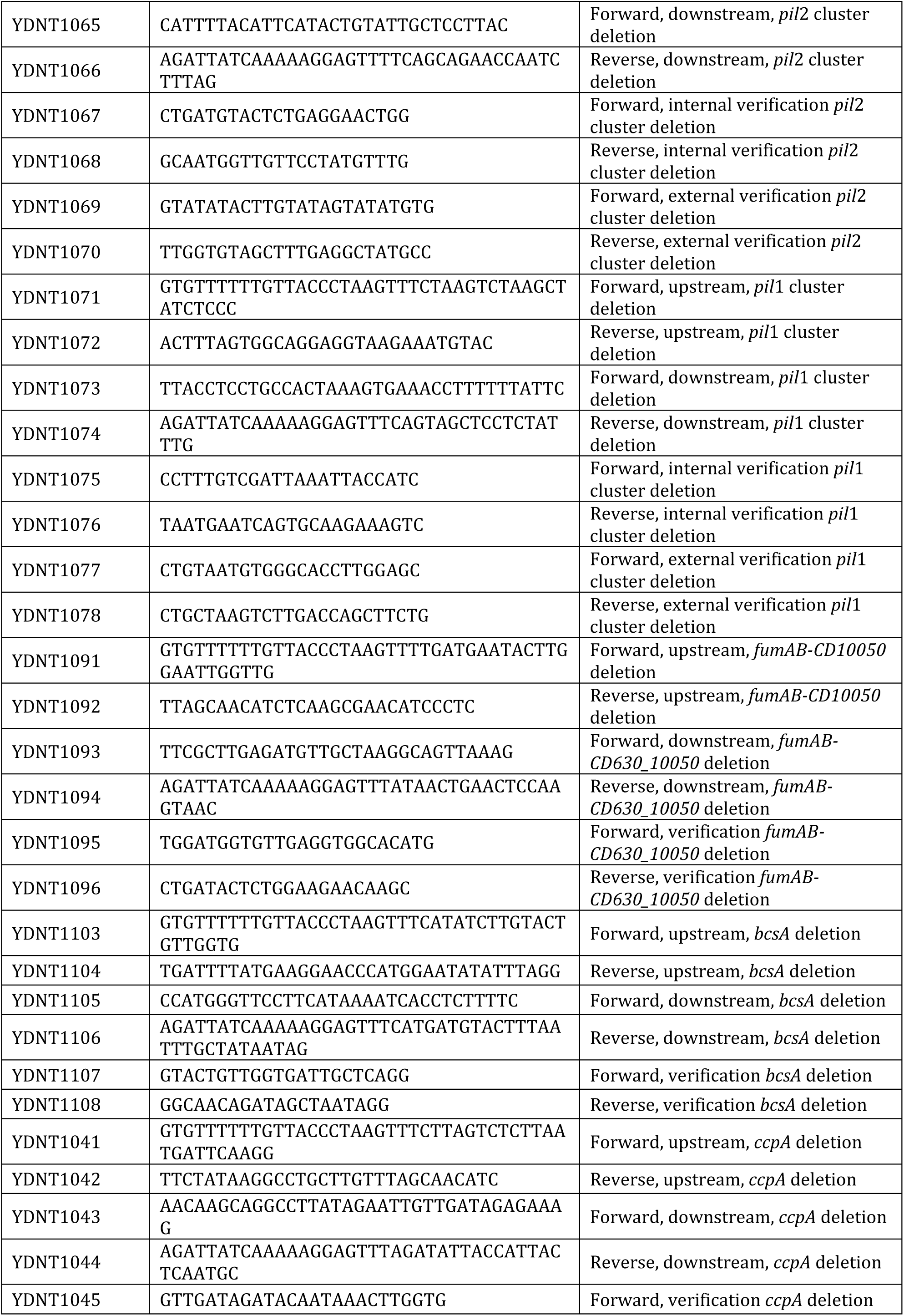
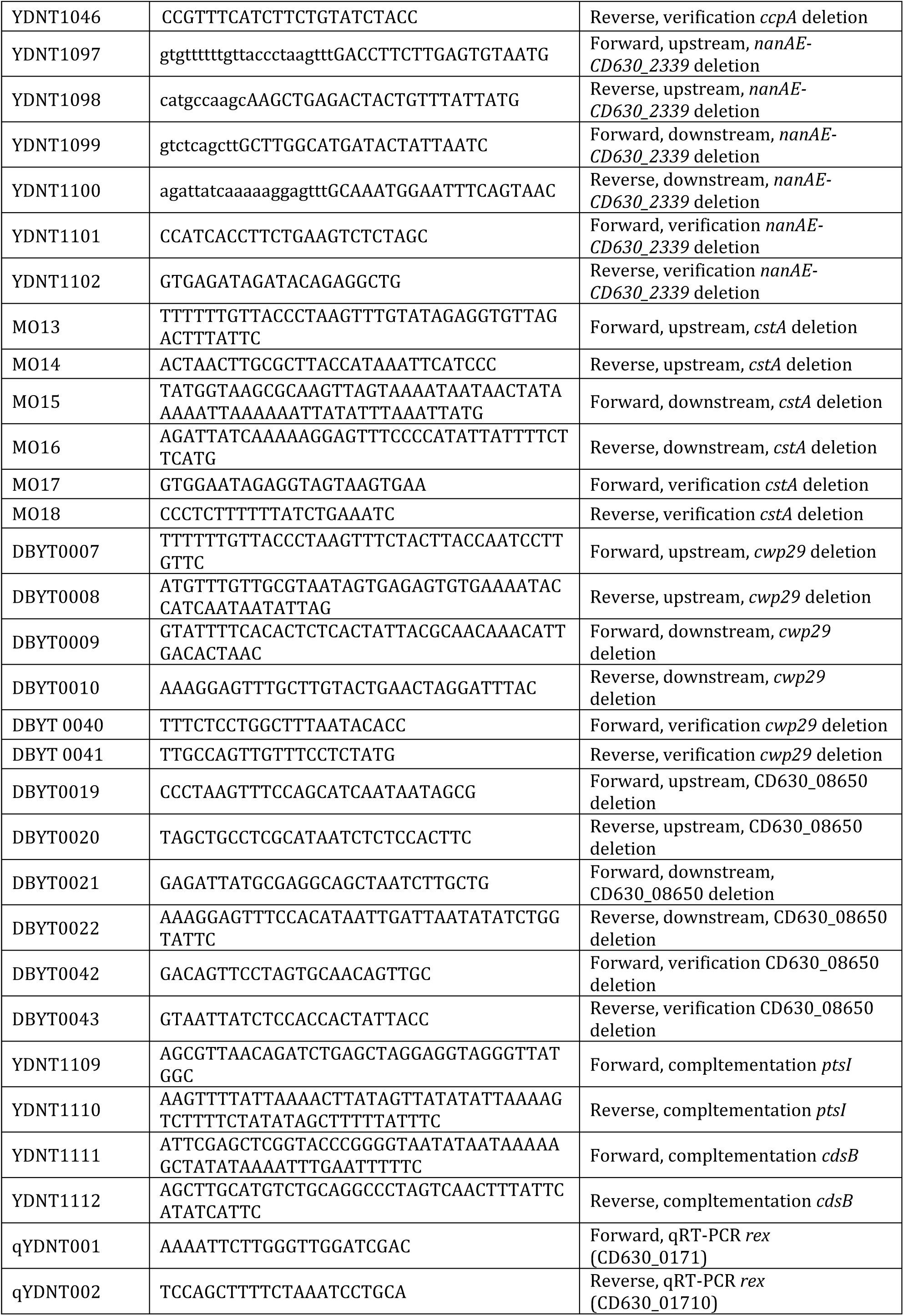
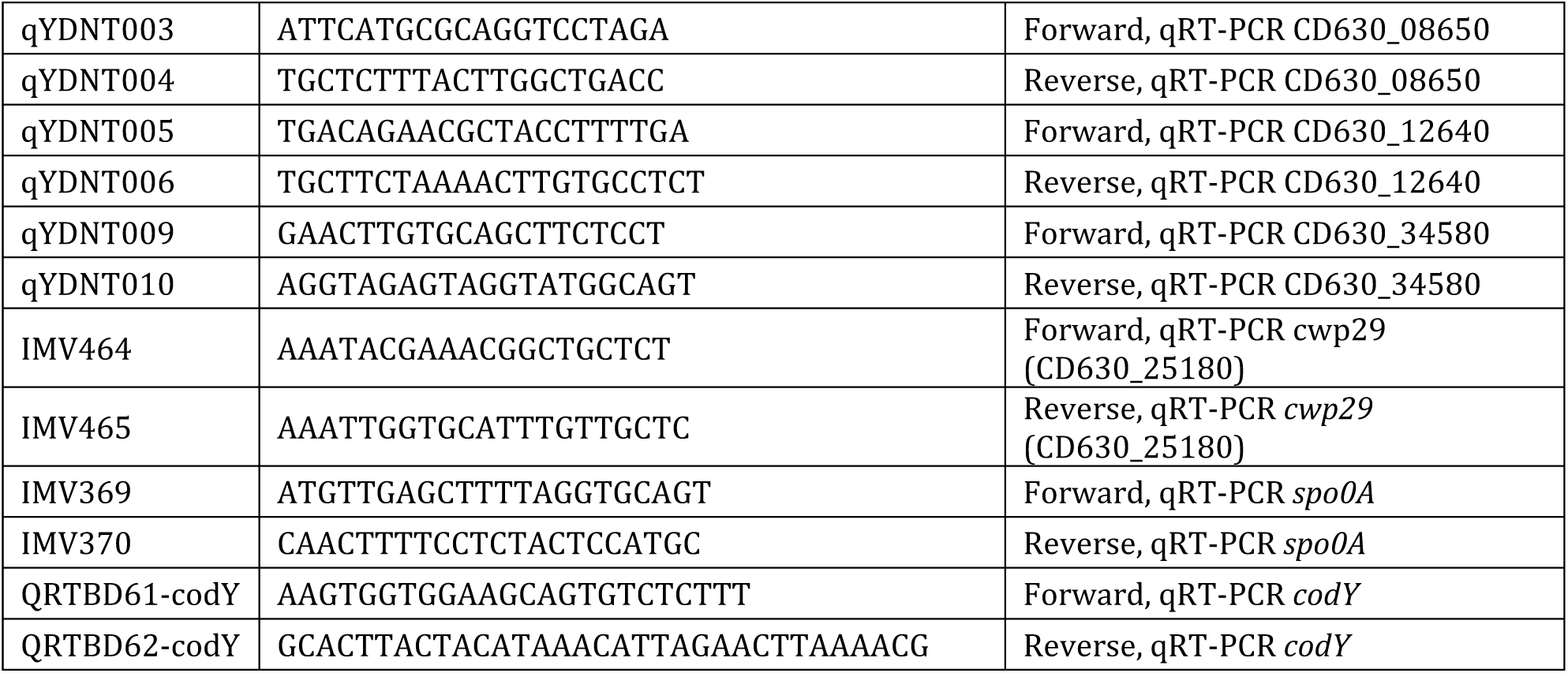
List of primers used for gene deletion and complementation constructs

